# Successful strategies in the voluntarily repeated Prisoner’s Dilemma

**DOI:** 10.64898/2026.01.16.699891

**Authors:** Luis R. Izquierdo, Segismundo S. Izquierdo, Robert Boyd

## Abstract

Reciprocity is central to explanations of cooperation among unrelated individuals in societies of humans and other animals. Most mathematical analyses of the evolution of reciprocity are based on the repeated Prisoner’s Dilemma (RPD) and typically assume that new strategies are rarely introduced by mutation or analogous cultural processes, that behavioral errors are absent or infrequent, and that agents are bound to interact with the same partner. Here we analyze a version of the RPD in which new strategies are frequently introduced, behavioral errors occur at a substantial rate, and actors may have the option to leave their current partner. In this environment, the usual indeterminacy disappears and the mix of strategies and cooperation levels are quite stable. With the option to leave, cooperation persists at a substantially higher level than without the option to leave. Classical strategies such as *Grim, Tit-for-Tat*, or *Win-Stay-Lose-Shift* disappear and are replaced by strategies that sanction cheaters by leaving rather than by retaliatory defection. Beyond a threshold, increasing the number of times partners interact decreases the level of cooperation without the option to leave, but increases it when leaving is possible.

**Significance Statement:** Cooperation is central to human societies, and the repeated Prisoner’s Dilemma has long been used to explain reciprocity. Most models assume that individuals must interact repeatedly with the same partner. We show that when behavioral errors and variability are substantial, allowing individuals to leave their partner fundamentally changes which strategies succeed and how much cooperation is sustained. Classical strategies such as *Grim, Tit-for-Tat*, and *Win-Stay-Lose-Shift* are eliminated. Instead, strategies that sanction cheaters by leaving prevail. Leaving both protects cooperators from exploitation and generates positive assortment, as similar strategies interact more often with each other. These results reveal a simple principle: freedom to leave can promote cooperation at the population level.

---

Cooperation among unrelated individuals is a defining feature of human societies and a central puzzle across the biological and social sciences. The repeated Prisoner’s Dilemma (RPD) has long served as the workhorse model to explain how pairwise reciprocity can emerge—by conditioning future actions on past behavior, individuals can sustain mutually beneficial cooperation.

The theoretical study of the RPD has largely been guided by an equilibrium-based perspective, in which predictions are framed in terms of equilibrium concepts such as Nash equilibrium and their refinements. A central difficulty in this approach is indeterminacy. For sufficiently large continuation probability, the Folk Theorem implies the existence of infinitely many Nash equilibria in the RPD, but none of them are strict. From an evolutionary perspective, there are infinitely many neutrally stable strategies (1, 2), but none are evolutionarily stable (3) or dynamically stable (4). Allowing for behavioral errors—so that players may occasionally choose unintended actions—alters this picture but does not eliminate the indeterminacy. With small errors, there can be a large number of strict Nash strategies (5–8), many of which can support full cooperation. These equilibria correspond to asymptotically stable states under most evolutionary dynamics.

One approach to obtaining sharper predictions has been to study evolutionary models with vanishingly small mutation rates (4, 9–20). In this regime, populations spend most of their time in homogeneous states, with evolutionary change occurring through rare transitions between them as mutants either go extinct or fix before further mutations arise. Long-run behavior is then characterized by the relative prevalence of equilibria, reinforcing the equilibrium-selection perspective that underlies much of the theoretical analysis of the RPD.

A central limitation of this body of work is the focus on settings with negligible or nonexistent errors and mutation. Empirical evidence from both biological and social systems indicates that behavioral errors and mutation are common, and that substantial variation is often maintained within populations (21–24). Importantly, previous work has shown that evolutionary dynamics with non-negligible mutation rates behave qualitatively differently from the rare-mutation limit (24–26). Substantial mutation and errors lead to distinct dynamical regimes rather than to small perturbations of equilibrium dynamics. This is the setting we consider here: an environment in which errors and mutation occur with appreciable probability.

A striking consequence of allowing substantial mutation and errors is that predictions often become sharper. When mutation and error rates are large enough to maintain persistent variation, the profusion of equilibria characteristic of repeated games collapses to a small number of dynamical attractors (25). This collapse makes it possible to move beyond equilibrium enumeration and instead identify the behavioral traits that characterize successful strategies, as well as the levels of cooperation sustained in the long run.

In addition to the standard RPD, we consider the voluntarily repeated Prisoner’s Dilemma (VRPD), in which players may terminate partnerships and seek a new partner (27–39). This setting captures situations in which individuals can leave uncooperative partners, such as friendships, marriages, economic relationships, and mutual aid in humans and other animals (40). Equilibrium analysis of the VRPD is even less selective than in the RPD: a wide range of partially cooperative equilibria exists (39), and evolutionary models with rare mutation likewise fail to identify a small set of successful strategies (37). As in the RPD, equilibrium multiplicity persists even in the presence of partner choice, and neither equilibrium analysis nor evolutionary models with rare mutation are sufficient to explain which behaviors may be observed.

Here we compare the RPD and the VRPD in an evolutionary framework with substantial behavioral errors and mutation. We focus on deterministic memory-one strategies (26), which includes the most widely studied strategies in the literature while avoiding unbounded strategic complexity. We assume that errors occur frequently and that mutation maintains significant variation in the population. In this setting, evolutionary trajectories in both games converge to a small number of robust regimes characterized by stable cooperation levels and well-defined behavioral traits. This allows us to identify which strategies are successful and to compare cooperation with and without the option to leave.

We find that, despite the absence of fully cooperative equilibria in the VRPD, cooperation is higher and more robust in the VRPD than in the RPD due to positive assortment generated by partner switching. This result aligns with experimental evidence (34, 41, 42), which shows that cooperation is higher when individuals can leave uncooperative partners and that partner switching is frequent when costs are not excessive. Our results also show that classical strategies—such as Tit-for-Tat (*TFT*) and Win-Stay-Lose-Shift (*WSLS*)—that have been celebrated for their success in the RPD (25, 43–45) perform poorly in the VRPD: with moderate error and mutation rates, these strategies are eliminated from the population. More broadly, our results add to growing evidence that analyses assuming zero error and vanishing mutation fail to capture the behavior of repeated interactions when variation is substantial.

## The model

Our model encompasses the standard RPD and the VRPD. The standard RPD is obtained as a special case of the VRPD by forbidding strategies that use the option to leave. Figure 1 shows a sketch of the sequence of events that take place in the model within each (discrete) time step (or tick). At the beginning of every tick, all single players (those without a partner) are randomly matched in partnerships, and then all agents play the stage game. Analysis presented in the SI indicates that our results are not substantially affected by delayed re-matching.

**Fig. 1.**
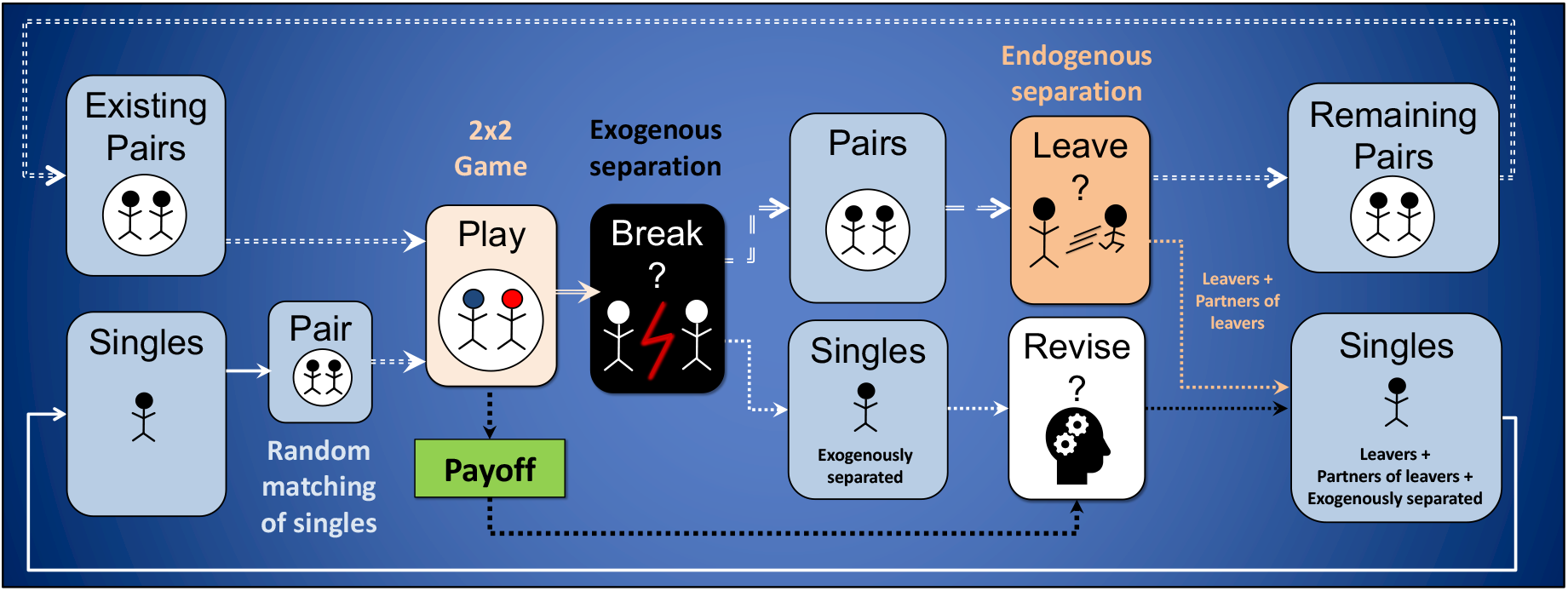
Sketch of the sequence of events within each time step. The “Remaining Pairs” and “Singles” at the end of a time step are identical to the “Existing Pairs” and “Singles” in the next time step.

The stage game is a Prisoner’s Dilemma (PD). Actors can either cooperate (*C*) or defect (*D*) with payoffs *U*_*CC*_ = *b, U*_*CD*_ = 0, *U*_*DC*_ = *b* + 1 and *U*_*DD*_ = 1 where the first action is the actor’s and the second their partner’s. This version of the PD is often called the donation game, where *b >* 1 is the benefit provided by a cooperating partner at a cost equal to 1. Each player’s strategy determines their action given the partnership’s history. We consider memory-one pure strategies, i.e., deterministic strategies whose choices depend only on the partnership’s last outcome. We can represent any of these strategies by a sequence of five letters indicating what choice to take (*C, D* or *L*, where *L* denotes leaving the partnership) after each possible history (*∅, CC, CD, DC, DD*). For example, *C*| *CLCC* is the strategy that starts a new partnership playing *C*, never plays *D*, and leaves after a partner’s unilateral defection. There are 2 *×* 3^4^ = 162 memory-one strategies in the VRPD. The memory-one strategies for a RPD are the 2*×* 2^4^ = 32 strategies with no *L*. For instance, *Tit-For-Tat* (*TFT*) corresponds to *C* |*CDCD*, while *C* |*CDDD* is a strategy that chooses to cooperate with new partners and continues to choose to cooperate if and only if there is mutual cooperation, defecting otherwise. In a slight abuse of convention, we call this last strategy *Grim*.

Errors are modeled as unintended actions in the stage game, a *D* instead of an intended *C*, or a *C* for a *D*. Both errors occur with probability *E*. After playing the stage game, partnerships may dissolve for exogenous reasons. The probability that this does not occur and the partnership continues is *δ*. Thus, *γ* = (1 − *δ*)^−1^ is the *expected number of interactions* of partnerships whose members do not choose to break up. After every exogenous breakup, players may revise their strategy independently, with probability *r*. Revisers experiment with probability *µ*, which we refer to as the *mutation* probability to maintain consistency with the evolutionary game theory literature, and imitate with probability 1 − *µ*. Experimentation introduces random behavioral variation through small changes in strategy (see SI for details), while imitation follows a payoff-based rule, whereby an agent is selected with probability proportional to payoffs and its strategy is copied. This updating process can be interpreted as cultural imitation, migration, or reproduction with mutation. Simulations were run in finite populations of size *N* ∈ [100, 1000] for sufficiently long durations to ensure that observed dynamics are representative of the long-run behavior of the underlying stochastic process. We also used a deterministic mean-dynamic approximation to study behavior in large populations (see SI).

## Simulation results

In this section we compare the RPD with the VRPD using computer simulation. As a baseline scenario, we consider: *b* = 3, *δ* = 0.95, *E* = 0.05, *µ* = 0.05, *r* = 0.1, and *N* = 500. In this setting, the option to leave increases cooperation from <40% to >60%, and this difference persists across a wide range of parameter values (Figure 2).

**Fig. 2.**
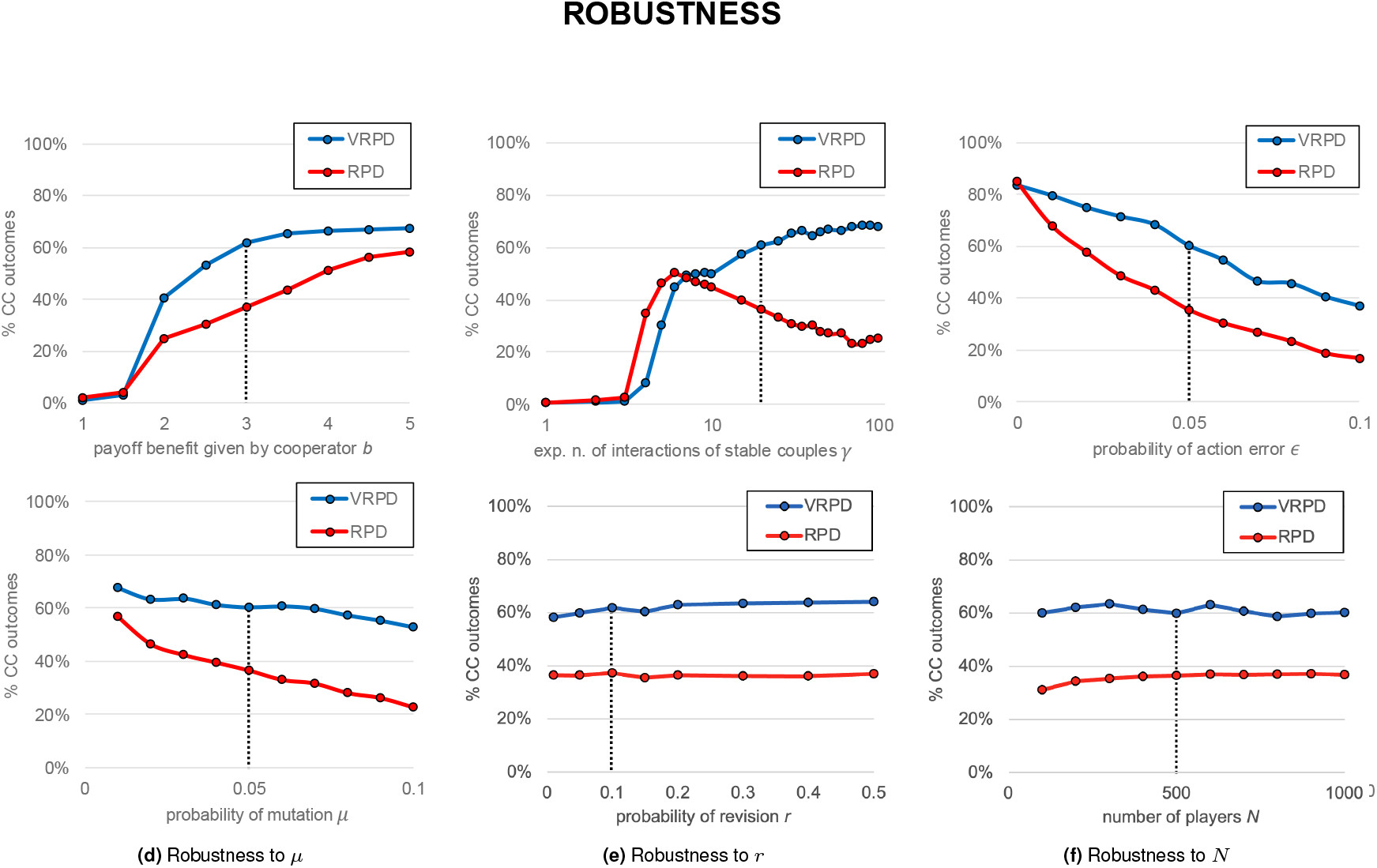
Average percentage of cooperative outcomes in the RPD and VRPD, departing from the baseline scenario, for different values of (a) the payoff benefit *b*, (b) the expected number of interactions of stable couples *γ*, (c) the probability of action error, (d) the probability of mutation *µ*, (e) the probability of revision *r*, and (f) the population size *N*. Each point is the average from tick 190 000 to tick 200 000 over 100 runs starting from random initial conditions. The standard error of every point is below 3%. The dashed lines indicate the values of the different parameters in the baseline scenario.

Figure 2(b) shows the appearance of cooperation at low values of *γ* in a logarithmic scale. If *γ* ≤ 3, cooperation is not observed: when partnerships last such a short time, the short-lived rewards enjoyed by partnerships of cooperators cannot compensate for exploitation by defectors. Cooperation begins to be significant at *γ* = 4. In the interval *γ* ∈ [4, 6], cooperation is significantly greater in the RPD than in the VRPD. Then, for larger values of *γ*, an interesting pattern appears: increasing the number of expected interactions (*γ*) lowers cooperation in the RPD, but increases cooperation in the VRPD. Thus, the cooperation gap between the RPD and VRPD widens with the continuation probability.

To understand this, we examine which strategies prevail in the baseline scenario for the RPD and for the VRPD. Table 1 shows that in the RPD selection favors: (1) being nice (87%), (2) cooperating after mutual cooperation (91%), (3) defecting after exploitation (91%), and (4) continuing to defect after exploiting (82%). The only two (memory-one) strategies with these four traits are *Grim* and *WSLS*. The difference between these two strategies is that *Grim* defects after mutual defection, while *WSLS* cooperates.

**Table 1.** Percentage of players using each choice (row) after a given history (column) in the RPD, in the baseline scenario. Values are averages from tick 290 000 to tick 300 000 over 100 runs starting from random initial conditions.

|  |  | History |  |  |  |  |
| --- | --- | --- | --- | --- | --- | --- |
| | | ∅ | CC | CD | DC | DD |
| Choice | C | 87 | 91 | 9 | 18 | 38 |
|  | D | 13 | 9 | 91 | 82 | 62 |

Figure 3 shows the four most prevalent strategies in the RPD simulations at the baseline scenario. They are all nice, keep on cooperating after mutual cooperation, and defect after having been exploited, i.e., they present the pattern *C* |*CD*○ ○ (where may ○ be *C* or *D*). In the absence of errors, any match between these four strategies leads to full cooperation.

**Fig. 3.**
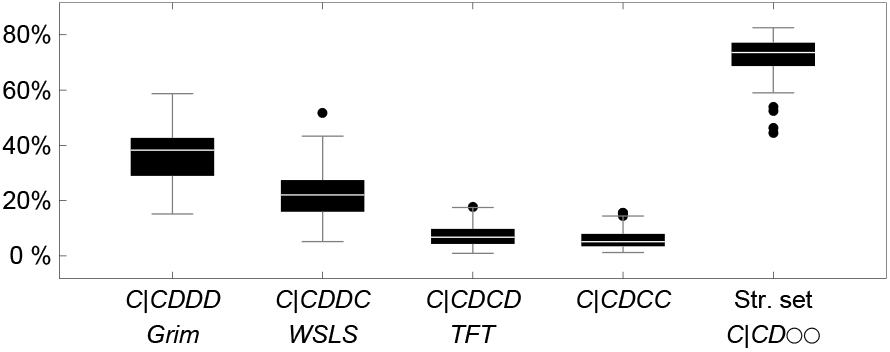
Box plots of the average usage of the four most successful strategies in the RPD, compiled over 100 runs starting from random initial conditions. Each average has been computed from tick 290 000 to tick 300 000. *C*|*CD*○ ○ represents the set of strategies where ○ may be *C* or *D*.

The presence of errors is crucial for understanding why increasing the expected number of interactions *γ* can lead to less cooperation (figure 2(b)). In the RPD, the only way cooperative strategies can protect themselves from exploiters is by defecting, and selection favors this trait: players defect after *CD* (see table 1). However, given this trait and with non-negligible errors, the longer a partnership lasts, the more likely it is that there will be mistakes that lead to sequences of mutual defection or miscoordinations. Thus, in the RPD, intermediate partnership lengths lead to the highest cooperation rates: if error rates are non-negligible the expected number of interactions should be neither too low nor too high.

Interestingly, while the presence of the strategy set *C* |*CD*○ ○ in the RPD is very significant and stable (see the right-most box plot in figure 3), there are strong fluctuations over time and between runs in the usage of each individual strategy in the set (see figure S1).

Now consider the VRPD. The possibility of leaving a partner introduces two important features. First, players can use the *defect-and-leave* strategy *D* |*LLLL*. In a population of nice strategies, *D* |*LLLL* players receive the maximum payoff *b* on every turn. This means that to be stable a population must have enough strategies that defect on the first period. Otherwise the population would be easily invaded by *defectand-leave*. In particular, *nice* strategies that cooperate with strangers (*TFT, WSLS, Grim*) can easily be invaded. Second, after having been exploited (possibly by mistake), players can now leave, rather than stay and defect, and this allows them to escape the potentially long sequence of mutual defections that strategies like *Grim* may suffer after a single mistake. The option to leave creates a different way to discipline defectors: sending them to the pool of singles by breaking the partnership. This punishment is more effective as the proportion of nasty strategies in the pool increases; and we know that there are some nasty strategies in any stable population, because nice strategies do not constitute Nash equilibria. With these two features in mind, it may not come as a surprise that the most successful strategies in the VRPD (table 2): (1) are *not* nice (97%), (2) continue cooperation after *CC* (89%), (3) leave after exploitation (82%)—this protects against defectors and generates positive assortment, and (4) cooperate after mutual defection (94%). We saw this last feature in *WSLS*, but here it exists for a different reason. In the RPD, *WSLS* cooperates after defection in order to restore cooperation after an error. Successful strategies in the VRPD cooperate after mutual defection to restore cooperation after the initial defections.

**Table 2.** Percentage of players using each choice (row) after a given history (column) in the VRPD. Values gathered as in table 1.

| Choice |  | History |  |  |  |  |
| --- | --- | --- | --- | --- | --- | --- |
| | | ∅ | $CC$ | $CD$ | $DC$ | $DD$ |
| | $C$ | 3 | 89 | 4 | 26 | 94 |
| | $D$ | 97 | 9 | 15 | 46 | 5 |
| | $L$ | | 2 | 82 | 28 | 1 |

Figure 4 shows the three most common strategies in the VRPD at the baseline scenario. These strategies, which form the set *D* |*CL* ●*C* (where may ● be *C, D* or *L*), are the only three strategies that present the four successful traits outlined above. They differ only in their behavior after exploiting their partner (i.e., after *DC*). After their initial defection, Leaver_1_ (*D* |*CLDC*) and Leaver_2_ (*D*| *CLLC*) behave similarly to *WSLS*, except that they respond to a partner’s defection by leaving rather than by defecting. Leaver_3_ (*D* |*CLCC*) cooperates after defecting. Given that 82% of the population leaves after having been exploited, behavior in this circumstance has only small effects on fitness. In the absence of errors, any match between these three strategies leads to the exact same path: *DD*-*CCCC*-*CC*-… However, if an error occurs the partnership is broken.

**Fig. 4.**
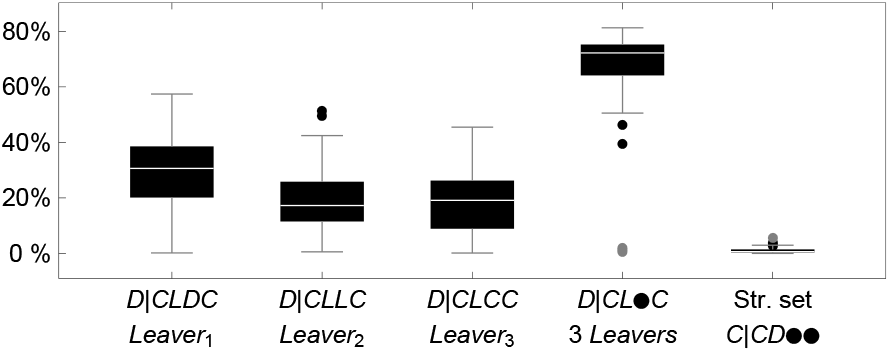
Box plots of the average usage of the three most successful strategies in the VRPD. Symbol ● in the definition of the strategy sets may be *C, D* or *L*. Values gathered as in figure 3.

The most successful strategies in the RPD, i.e. *C* |*CD○ ○* (72%), disappear when players are given the option to leave (see the right-most box plot in figure 4). Instead, it is the set *D* |*CL ●C*, that persists as a significant fraction (67%) of the population. And by exercising the option to leave, these strategies produce a higher cooperation rate (from less than 40% to more than 60% in the baseline scenario; see figure 2). Like in the RPD, in the VRPD there are also strong fluctuations in the usage of successful strategies (figure S2).

Finally, note that, in the VRPD, increasing the expected number of interactions, *γ*, does not lead to less cooperation, as happens in the RPD (see figure 2(b)). The reason is that successful players do not stay and defect after having been exploited; they simply leave and so escape the potentially long sequence of mutual defections that classical strategies may suffer after a single mistake. Strategies that do not leave suffer a difficult dilemma: if they are not tolerant of errors, like *Grim*, then they do not fare well against themselves; if, on the other hand, they are tolerant of errors, like *WSLS*, then they are vulnerable to exploitation by defectors. Leaving provides a way out of this dilemma: if you do fairly well against players using your strategy, then you can protect yourself from defectors by leaving them, and this generates positive assortment, a key principle known to underlie the evolution of cooperation (46–48). The overall level of cooperation is greatly increased in comparison with a system where players are not allowed to leave their partners. More freedom at the individual level leads to better outcomes at the population level.

## Mathematical insights

The simulation results have been confirmed and extended with analytical work described in the SI. There, we formalize important differences between the RPD and the VRPD in the presence of errors. In the RPD with errors there are a number of monomorphic strict Nash states to which many evolutionary dynamics can converge, but that is not the case in the VRPD. For instance, in our baseline scenario, *Grim, WSLS, AllD*, and *D CDDC* (*nWSLS*, for *nasty WSLS*) are strict Nash in the RPD, but they are not even evolutionarily stable in the VRPD. In fact, focusing on the donation game, we show that no memory-one strategy is evolutionarily stable (or strict Nash) in the VRPD for moderate error rates (*ϵ* ∈ (0, 0.1]) and *δ* ∈ [0.5, 1).

In the SI we also derive the mean dynamic (MD) of our model. The MD (49) is a deterministic system of ODEs that approximates the (stochastic) dynamics of the population over finite time spans. We solved it numerically to identify attractors and better understand the success of strategies that exercise the option to leave.

In particular, we explore the performance of *Leaver*_1_ in a population initially composed of the three most successful strategies in the RPD (i.e., *Grim, WSLS*, and *TFT*) and *AllD*. We numerically solve this MD starting from 1000 random initial conditions with baseline parameters and find that all trajectories converge to a state where more than 80% of players are using *Leaver*_1_. Figure 5 shows how the strategy distribution of the seemingly unique global attractor of this MD changes with the mutation rate *µ*. As the mutation rate *µ* tends to zero, *Leaver*_1_ increases its dominance and ends up completely eliminating the four strategies that do not leave.

**Fig. 5.**
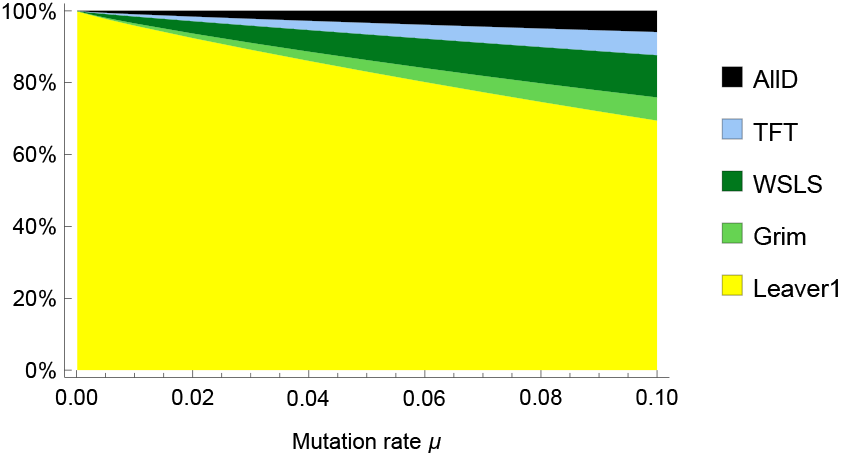
Strategy distribution of the seemingly unique global attractor of the MD for different mutation rates. Other parameter values are set as in the baseline scenario. This attractor is not necessarily stable with a larger strategy set.

We also study the attractors of this reduced system for different error rates (see figure 6). For *ϵ* ≤10^−3^ there seems to be a global attractor where more than 75% of the population is using *TFT, WSLS* or *Grim*. Then there is a small region of values of *E* for which there is more than one attractor.

**Fig. 6.**
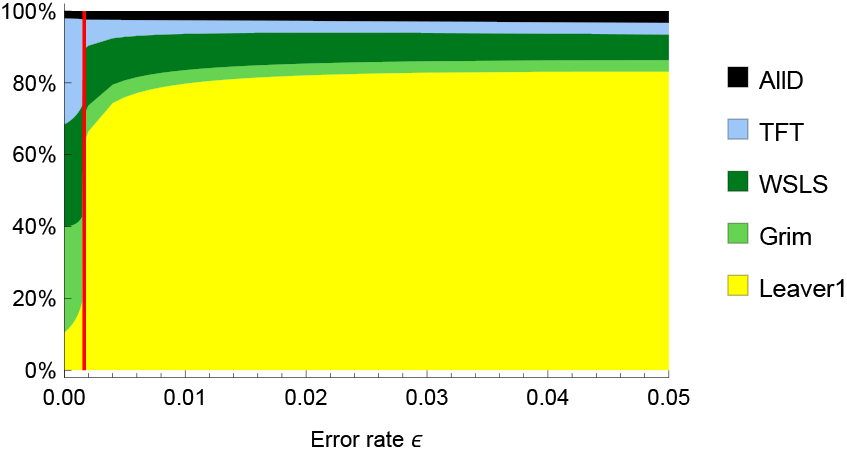
Strategy distribution of the seemingly unique global attractor of the MD for different values of the error rate ϵ∈ [0, 0.05]. Other parameter values are set as in the baseline scenario. For each value of, the MD is numerically solved starting from each of the 5 corners of the simplex. In the region colored in red, trajectories starting from different corners converge to two different distributions, so there is no global attractor in that region. For all other values of ϵ, the 5 initial conditions converged to the distribution shown in the plot.

And finally, for ϵ ≥ 2 *×*10^−3^, a new global attractor appears, completely different from the one present for *ϵ ≤* 10^−3^. This new attractor is characterized by the dominance of *Leaver*_1_, and stays for greater values of *ϵ*.

Figure 6 shows that the dynamics of these models for small error rates, e.g. *ϵ*= 2 *×*10^−3^, can be completely different from the dynamics where the error rate is infinitesimally small. Thus, conclusions based on limits where the error rate tends to zero, or is actually zero, are not necessarily valid for systems where the error rate is very low, but not vanishingly small.

## Leaving vs punishing by defecting. Positive assortment

A comparison of *Leaver*_1_ (*D*| *CLDC*) and *nWSLS* (*D* |*CDDC*) illustrates why leaving unilateral defectors is better than punishing them by defecting (like *nWSLS*).

First, notice that in a population made up of any mixture of *Leaver*_1_ and *nWSLS* both strategies obtain the same expected payoff (see figure 7(a)) because in every partnership individuals expect the same sequence of outcomes. Without errors, the sequence of outcomes in every partnership is *DD*-*CC*-*CC*-… and after an error the expected sequence is also *DD*-*CC*-*CC*-… Figure 7(a) includes the (1-dimensional) mean dynamic at the bottom: arrows point in the direction of movement and circles indicate the location of the asymptotically stable rest points.

**Fig. 7.**
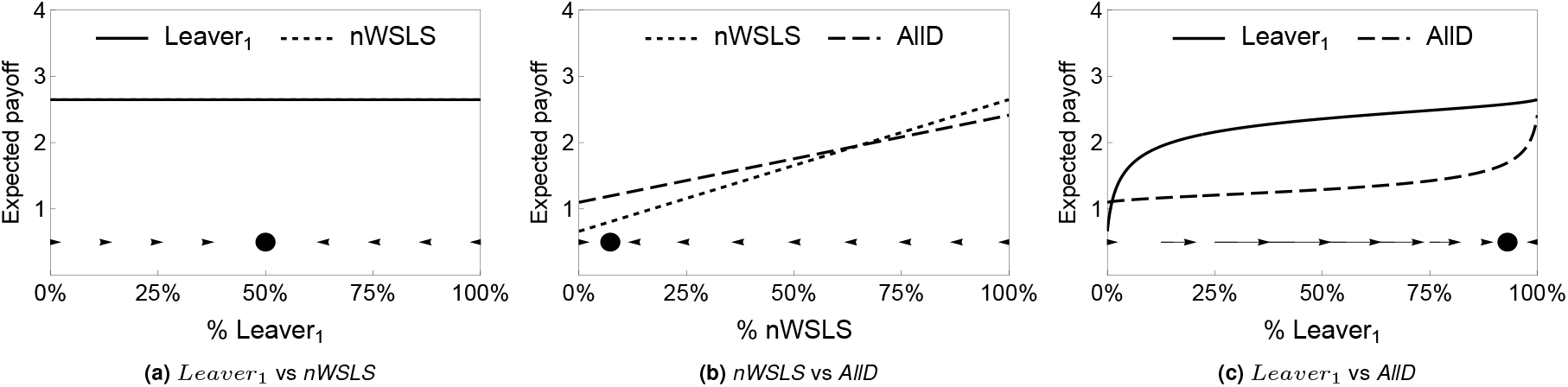
Payoff functions and phase portraits for several bimorphic populations. Parameters are set as in the baseline scenario (*δ* = 0.95, and = *µ* = 0.05). Mutation in these figures consists in choosing one of the two strategies at random. Note that errors affect the expected payoff obtained by each strategy, but mutation does not.

Next, consider the fitness for *nWSLS* and *Leaver*_1_ interacting with *AllD* (*D* |*DDDD*, see figures 7(b) and 7(c) respectively). An individual with either strategy obtains the same payoff in a population of *AllD*, and a single *AllD* obtains the same payoff against either of them when entering a monomorphic population. In both cases a player using *AllD* can expect an alternating sequence of *DD* and *DC* outcomes (either with the same partner if in a *nWSLS* population, or changing partners after *DC* if in a *Leaver*_1_ population).

However, the fitnesses of *nWSLS* and *Leaver*_1_ differ in populations with substantial frequency of *AllD* as shown in figure 7. Panel 7(a) shows that both strategies are equivalent in an isolated bimorphic population. Panel 7(b) shows the dynamics in a population of *nWSLS* and *AllD*: linear payoff functions and a single attractor where most players use *AllD*. Panel 7(c) shows the dynamics in a population of *Leaver*_1_ and *AllD*: nonlinear payoff functions (with the same initial and final values as in panel 7(b)) and a single attractor where most players use *Leaver*_1_.

To understand why *nWSLS* and *Leaver*_1_ differ, assume that the frequency of *AllD* defectors is 10%. The first part of Table 3 presents the expected fraction of *nWSLS* players in partnership with *nWSLS* or with *AllD* players. Since partnerships are never endogenously broken, those are simply the proportions of each strategy in the population, and the same holds for *AllD* players. The second part of the table shows the average payoff obtained by *nWSLS* and *AllD* players in partnerships with either *nWSLS* or *AllD*. The last column shows the average payoff obtained by each strategy in the population.

**Table 3.** Assortment and payoffs in a bimorphic population with 90% of *nWSLS* players and 10% of *AllD* players. *δ* = 0.95 and *ϵ* = 0.05.

|  | Fraction matched to |  | Payoff with |  | Payoff |
| --- | --- | --- | --- | --- | --- |
|  | *nWSLS* (90%) | *AIID* (10%) | *nWSLS* | *AIID* |  |
| *nWSLS* | 90% | 10% | 2.65 | 0.66 | 2.45 |
| *AIID* | 90% | 10% | 2.41 | 1.10 | 2.28 |

Table 4 presents the corresponding data for *Leaver*_1_ and *AllD*. In this case, homogeneous partnerships in which both players use the same strategy are stable, while heterogeneous partnerships separate after playing together twice (unless there is an error). As a result, most *Leaver*_1_ players are found in partnerships with other *Leaver*_1_ players, and most *AllD* players are found in partnerships with other *AllD* players. The option to leave generates positive assortment: the fraction of *Leaver*_1_ players who are in partnerships with other *Leaver*_1_ players is larger than the proportion of *Leaver*_1_ players in the population; and the same holds for *AllD* players. As a result of positive assortment, the average payoff to *Leaver*_1_ players in the population is close to the payoff obtained when two such players interact (this effect *pushes* its payoff function upwards in figure 7(c) compared to 7(b)); and the same holds for *AllD* players (which *pushes* its payoff function downwards in figure 7(c)).

**Table 4.** Assortment and payoffs in a bimorphic population with 90% of *Leaver*_1_ players and 10% of *AllD* players. *δ* = 0.95 and *ϵ* = 0.05.

|  | Fraction matched to |  | Payoff with |  | Payoff |
| --- | --- | --- | --- | --- | --- |
|  | *Leaver*<sub>1</sub> (90%) | *AIID* (10%) | *Leaver*<sub>1</sub> | *AIID* |  |
| *Leaver*<sub>1</sub> | 95.65% | 4.35% | 2.65 | 0.66 | 2.56 |
| *AIID* | 39.14% | 60.86% | 2.41 | 1.10 | 1.61 |

## Conclusions

We have studied population dynamics in the repeated Prisoner’s Dilemma and in the voluntarily repeated Prisoner’s Dilemma with the possibility of execution errors and with frequent exploration of alternative strategies. Both features are commonly observed in real-life interactions but have often been considered separately. Studying them together leads to insights not captured by previous models.

In settings with significant errors and mutation, a small set of winning strategies typically survives. While precluding a fully cooperative equilibrium, the option to leave significantly enhances cooperation, unless the expected number of interactions is very small (see also (50)). This boost in cooperation increases with the number of expected interactions. The set of surviving strategies varies completely with the option to leave. Strategies that exercise the option to leave are more robust to errors and thrive compared to strategies that never leave. With errors, non-leaving strategies such as *Grim, Win-Stay-Lose-Shift* or *Tit-for-Tat* face a dilemma: forgiveness invites exploitation, while strictness causes long miscoordination. Leaving resolves this by letting players abandon exploiters while still cooperating with similar types. As a result, strategies that leave unilateral defectors outperform those who punish by defecting within partnerships, and cooperation increases markedly.

Importantly, the option to leave not only provides an effective alternative to in-partnership retaliation. It also generates positive assortment, allowing cooperative strategies who leave defectors to interact more often with each other. Cooperative partnerships persist, while defectors often return to the pool of singles, which is dominated by defectors. Thus, most cooperators are found in partnerships with other cooperators, and most defectors are found in partnerships with defectors. This allows for the evolutionary emergence and maintenance of high levels of cooperation.

In sum, our results highlight the central role of the option to leave in supporting (partial) cooperation under realistic conditions. Giving agents freedom to leave exploitative relationships can lead to better outcomes at the population level.

## ACKNOWLEDGMENTS

Financial support from the Spanish State Research Agency (PID2024-159461NB-I00/MICIU and PID2020-118906GB-I00/MCIN), Junta de Castilla y León and EUFEDER program (CLU-2025-2-05 - BIOECOUVA), and the Institute of Human Origins, Arizona State University is gratefully acknowledged.

## References

1. J Bendor and P Swistak. Types of evolutionary stability and the problem of cooperation. Proc. National Acad. Sci. United States America, 92(8):3596–3600, 04 1995.

2. Jonathan Bendor and Piotr Swistak. Evolutionary equilibria: Characterization theorems and their implications. Theory Decision, 45:99–159, 1998.

3. Robert Boyd and Jeffrey P. Lorberbaum. No pure strategy is evolutionarily stable in the repeated prisoner’s dilemma game. Nature, 327(6117):58–59, 1987.

4. Julián García and Matthijs van Veelen. In and out of equilibrium I: Evolution of strategies in repeated games with discounting. J. Econ. Theory, 161:161 – 189, 2016.

5. Robert Sugden. The economics of rights, co-operation and welfare. Basil Blackwell, Oxford, 1986.

6. Robert Boyd. Mistakes allow evolutionary stability in the repeated prisoner’s dilemma game. J. Theor. Biol., 136:47–56, 1 1989.

7. Yong Gwan Kim. Evolutionarily stable strategies in the repeated prisoner’s dilemma. Math. Soc. Sci., 28:167–197, 12 1994.

8. Olof Leimar. Repeated games: A state space approach. J. Theor. Biol., 184(4):471–498, 1997.

9. Michihiro Kandori, George J. Mailath, and Rafael Rob. Learning, mutation, and long run equilibria in games. Econometrica, 61:29–56, 1993.

10. H. Peyton Young. The evolution of conventions. Econometrica, 61:57–84, 1993.

11. H. Peyton Young. Individual Strategy and Social Structure. Princeton University Press, Princeton, 1998.

12. Drew Fudenberg and Lorens A. Imhof. Imitation processes with small mutations. J. Econ. Theory, 131:251–262, 2006.

13. Drew Fudenberg, Martin A. Nowak, Christine Taylor, and Lorens A. Imhof. Evolutionary game dynamics in finite populations with strong selection and weak mutation. Theor. Popul. Biol., 70:352–363, 2006.

14. Arne Traulsen and Christoph Hauert. Stochastic evolutionary game dynamics. In EdHeinz Georg Schuster, editor, Reviews of Nonlinear Dynamics and Complexity, volume 2, pages 25–61. Wiley, New York, 2009.

15. Bin Wu, Chaitanya S. Gokhale, Long Wang, and Arne Traulsen. How small are small mutation rates? J. Math. Biol., 64(5):803–827, 2012.

16. Christian Hilbe, Martin A. Nowak, and Karl Sigmund. Evolution of extortion in iterated prisoner’s dilemma games. Proc. National Acad. Sci., 110(17):6913–6918, 2013.

17. Alexander J. Stewart and Joshua B. Plotkin. From extortion to generosity, evolution in the iterated prisoner’s dilemma. Proc. National Acad. Sci., 110(38):15348–15353, 2013.

18. Alexander J. Stewart, Todd L. Parsons, and Joshua B. Plotkin. Evolutionary consequences of behavioral diversity. Proc. National Acad. Sci., 113(45):E7003–E7009, 2016.

19. Matthijs van Veelen and Julián García. In and out of equilibrium ii: Evolution in repeated games with discounting and complexity costs. Games Econ. Behavior, 115:113–130, 2019.

20. Laura Schmid, Krishnendu Chatterjee, Christian Hilbe, and Martin A. Nowak. A unified framework of direct and indirect reciprocity. Nat. Hum. Behaviour, 5(10):1292–1302, 2021.

21. Arne Traulsen, Dirk Semmann, Ralf D. Sommerfeld, Hans-Jürgen Krambeck, and Manfred Milinski. Human strategy updating in evolutionary games. Proc. National Acad. Sci., 107:2962–2966, 2 2010.

22. Jelena Grujić, Carlos Gracia-Lázaro, Manfred Milinski, Dirk Semmann, Arne Traulsen, José A. Cuesta, Yamir Moreno, and Angel Sánchez. A comparative analysis of spatial prisoner’s dilemma experiments: Conditional cooperation and payoff irrelevance. Sci. Reports, 4:4615, 4 2014.

23. Robert Boyd and Sarah Mathew. Arbitration supports reciprocity when there are frequent perception errors. Nat. Hum. Behaviour, 5(5):596–603, 2021.

24. Josef Tkadlec, Christian Hilbe, and Martin A. Nowak. Mutation enhances cooperation in direct reciprocity. Proc. National Acad. Sci., 120:e2221080120, 5 2023.

25. Lorens A. Imhof, Drew Fudenberg, and Martin A. Nowak. Evolutionary cycles of cooperation and defection. Proc. National Acad. Sci., 102(31):10797–10800, 2005.

26. Christian Hilbe, Luis A. Martinez-Vaquero, Krishnendu Chatterjee, and Martin A. Nowak. Memory-n strategies of direct reciprocity. Proc. National Acad. Sci., 114(18):4715–4720, 2017.

27. Nahoko Hayashi and Toshio Yamagishi. Selective play: Choosing partners in an uncertain world. Pers. Soc. Psychol. Review, 2:276–289, 11 1998.

28. James W. Friedman and Peter Hammerstein. To trade, or not to trade; that is the question, pages 257–275. Springer Berlin Heidelberg, 1991.

29. Magnus Enquist and Olof Leimar. The evolution of cooperation in mobile organisms. Animal Behaviour, 45:747–757, 4 1993.

30. Ronald Noë and Peter Hammerstein. Biological markets. Trends Ecol. & Evol., 10:336–339, 8 1995.

31. C. Athena Aktipis. Know when to walk away: contingent movement and the evolution of cooperation. J. Theor. Biol., 231:249–260, 11 2004.

32. C. Athena Aktipis. Is cooperation viable in mobile organisms? simple walk away rule favors the evolution of cooperation in groups. Evol. Hum. Behavior, 32:263–276, 7 2011.

33. Takako Fujiwara-Greve and Masahiro Okuno-Fujiwara. Voluntarily separable repeated prisoner’s dilemma. Review Econ. Stud., 76(3):993–1021, 2009.

34. Pat Barclay and Nichola Raihani. Partner choice versus punishment in human prisoner’s dilemmas. Evol. Hum. Behavior, 37(4):263–271, 2016.

35. Segismundo S. Izquierdo, Luis R. Izquierdo, and Fernando Vega-Redondo. The option to leave: Conditional dissociation in the evolution of cooperation. J. Theor. Biol., 267(1):76– 84, 2010.

36. Pat Barclay. Strategies for cooperation in biological markets, especially for humans. Evol. Hum. Behavior, 34:164–175, 5 2013.

37. Christopher Graser, Takako Fujiwara-Greve, Julián García, and Matthijs van Veelen. Repeated games with partner choice. PLOS Comput. Biol., 21:e1012810, 2 2025.

38. Alejandro Gutiérrez-Mielgo, Luis Izquierdo Millán, and Segismundo Izquierdo Millán. Games with costly endogenous separation. In Proceedings of the 1st International Electronic Conference on Games, 2025.

39. Segismundo S. Izquierdo and Luis R. Izquierdo. Stable strategies in repeated games with endogenous separation. Universidad de Valladolid Discuss. Pap., 2026.

40. Sarah Mathew and Robert Boyd. Rethinking reciprocity. Trends Cogn. Sci., inpress.

41. Katrin Fehl, Daniel J. van der Post, and Dirk Semmann. Co-evolution of behaviour and social network structure promotes human cooperation. Ecol. Lett., 14:546–551, 6 2011.

42. David G. Rand, Samuel Arbesman, and Nicholas A. Christakis. Dynamic social networks promote cooperation in experiments with humans. Proc. National Acad. Sci., 108:19193– 19198, 11 2011.

43. Robert Axelrod. The Evolution of Cooperation. Basic Books, 1984.

44. David Kraines and Vivian Kraines. Learning to cooperate with pavlov an adaptive strategy for the iterated prisoner’s dilemma with noise. Theory Decision, 35(2):107–150, 1993.

45. Martin Nowak and Karl Sigmund. A strategy of win-stay, lose-shift that outperforms tit-for-tat in the prisoner’s dilemma game. Nature, 364(6432):56–58, 1993.

46. Ilan Eshel and L. L. Cavalli-Sforza. Assortment of encounters and evolution of cooperativeness. Proc. National Acad. Sci., 79(4):1331–1335, 1982.

47. Michael Doebeli and Christoph Hauert. Models of cooperation based on the prisoner’s dilemma and the snowdrift game. Ecol. Lett., 8(7):748–766, 2005.

48. Jeffrey A Fletcher and Michael Doebeli. A simple and general explanation for the evolution of altruism. Proc. Royal Soc. B: Biol. Sci., 276(1654):13–19, 2009.

49. William H. Sandholm. Population games and evolutionary dynamics. The MIT Press, 2010.

50. Matthias Wubs, Redouan Bshary, and Laurent Lehmann. Coevolution between positive reciprocity, punishment, and partner switching in repeated interactions. Proc. Royal Soc. B: Biol. Sci., 283(1832):20160488, 2016.

