## Supporting information for "Successful strategies in the voluntarily repeated Prisoner’s Dilemma"

#### This PDF file includes:

Supporting text
Figs. S1 to S8
Tables S1 to S2
SI References

### Software

All the software used in this paper (i.e., a *NetLogo* agent-based model and several *Mathematica* notebooks) has been released under GNU General Public License v3 and can be downloaded for free at https://doi.org/10.5281/zenodo.17473910. Instructions to run the agent-based model can be found at https://luis-r-izquierdo.github.io/ES-1-outcome/.

### Supporting Information Text

This supporting text presents a clarification on how the mutation mechanism works in the stochastic model (section A), reports the results of some computational experiments designed to explore the effect of delayed rematching (section B), and explains the details of our mathematical analysis. The mathematical analysis is structured in six parts. The first and most foundational part (section C) is the derivation of the expected payoff, $F_i^\epsilon(\boldsymbol{x})$, for any strategy $i$ in any population state $\boldsymbol{x}$ with an error rate $\epsilon$. This calculation is non-trivial as it must account for endogenous separation, which alters the composition of the pool of singles relative to the whole population. We solve for the stationary distributions of partnership histories and outcomes using matrix equations and formally link the strategy distribution in the pool ($\boldsymbol{p}$) to the overall population distribution ($\boldsymbol{x}$).

With the payoff function established, the second part (section D) uses it to investigate static stability concepts. We test for strict Nash and evolutionarily stable strategies (ESS) and prove that, unlike in the RPD with errors, no memory-one strategy is evolutionarily stable (or strict Nash) in the VRPD for moderate error rates ($\epsilon \in (0, 0.1]$) and $\delta \in [0.5, 1)$ on the donation game (Proposition 1).

Section E uses $F_i^\epsilon(\boldsymbol{x})$ to derive the mean dynamic (MD) of the system. This system of differential equations provides a deterministic approximation of the stochastic evolutionary dynamics, allowing us to numerically solve for the system's attractors. Section F uses the MD to explain why leavers dominate in the VRPD.

Section G includes a figure created to help understand the sharp change in the distribution of strategies shown in figure 6 of the paper. Finally, section H includes the proof of Proposition 1 and of other formal statements.

**A. Mutation (or experimentation) mechanism.** When an individual revises its strategy, mutation occurs with probability $\mu$. Mutation consists of switching to a randomly chosen "neighboring strategy", defined as a strategy whose representation (as a list of 5 letters) differs in exactly one action. Operationally, the revising agent selects one of the five letters that define its memory-one strategy with equal probability and replaces the corresponding action with an alternative one. When more than one alternative action is available, the new action is chosen uniformly at random. This local mutation rule ensures that exploration introduces small, incremental changes in behavior while maintaining a well-defined notion of strategic proximity.

Note also that in genetic models, mutation rates at single loci are typically much lower than we assume. However, complex behavioral strategies are affected by many loci with small effects, and substantial heritable genetic variation exists at equilibrium (1). High mutation rates are a crude way to modeling such variation (e.g. (2)). In social contexts, variation could be due to individual experimentation or from errors in strategy imitation. Evidence from social learning experiments and naturalistic observations suggests that such error rates are substantial (3–5).

**B. Delayed rematching.** In this section, we explore the effect of delayed rematching. The NetLogo model includes two parameters that are not discussed in the main body of the paper:

- ***ticks-until-rematching***: the number of ticks that single players must wait in the pool of singles before being randomly rematched.
- ***out-payoff***: the payoff obtained by single players who are not rematched in the current period. If *ticks-until-rematching* equals 0, no player receives this payoff.

Individuals who become single must wait for *ticks-until-rematching* time steps before being rematched. Any individual who has spent exactly *ticks-until-rematching* ticks in the pool of singles is randomly paired with another single individual. Thus, when *ticks-until-rematching* $= 0$ (the case analyzed in the main text) every single individual is paired at the beginning of each time step.

Figure S3 reports the average percentage of cooperative outcomes in the RPD and VRPD for values of *ticks-until-rematching* ranging from 0 (immediate rematching, as in the paper) to 10. Throughout these simulations, *out-payoff* is set to 0. Results for the RPD (where delayed rematching should have no effect) are included to illustrate the level of variability attributable solely to stochasticity. Overall, introducing delayed rematching has no significant impact on cooperation levels in the VRPD, at least for values of *ticks-until-rematching* up to 10.

Finally, Table S1 shows that although delayed rematching induces small changes in the distribution of successful behavioral traits, these differences remain minor for values of *ticks-until-rematching* up to 10.

**C. Payoffs in a repeated game with endogenous separation and errors.** In this section we show how to calculate the expected payoff $F_i^\epsilon(\boldsymbol{x})$ obtained by a player using strategy $i$ in a population of players with strategy distribution $\boldsymbol{x}$ when the error rate is $\epsilon$. The results are valid for any symmetric $2 \times 2$ stage game and can be easily generalized to stage games with any number of actions. The actions of the stage game are $\{C, D\}$ and the payoffs are $U_{CC}, U_{CD}$, $U_{DC}$ and $U_{DD}$. The (finite) set of memory-one strategies is $\Omega$, and a distribution of strategies $\boldsymbol{x}$ is a point in the simplex $\Delta_\Omega$, whose components add up to 1.

Let an $ij$ partnership be a partnership in which one player uses strategy $i$ and the other player uses strategy $j$. Let $p_i$ denote the proportion of $i$-players in the *pool of singles* (which will usually be different from the proportion of $i$-players in the population, $x_i$). At a steady state (where the distribution of strategies in the pool of singles is assumed to be stable), we can calculate, from the distribution of strategies $\boldsymbol{p} \in \Delta_\Omega$ in the pool of singles:

- The distribution of strategies $\boldsymbol{x}$ in the whole population.
- The distribution of $ij$ partnerships playing each action profile.

This will allow us to calculate, for a given population distribution $\boldsymbol{x}$, the corresponding pool distribution $\boldsymbol{p}$ and the expected payoff obtained by each strategy.

- To calculate the fraction of new partnerships, consider the step of the process at which players look at their history to choose an action for the stage game. For each possible history $h \in H \equiv \{\emptyset, CC, CD, DC, DD\}$, let $z_h^{ij\epsilon}$ be the fraction of $i$-players, out of those in an $ij$ partnership, whose history is $h$. $z_\emptyset^{ij\epsilon}$ is the fraction of new $ij$ partnerships, so the ratio of all $ij$ partnerships to new $ij$ partnerships is $(z_\emptyset^{ij\epsilon})^{-1}$.
- To calculate average payoffs, consider the step of the process at which players have just played a stage game. For each possible action profile $o \in O \equiv \{CC, CD, DC, DD\}$, let $w_o^{ij\epsilon}$ be the fraction of $i$-players, out of those in an $ij$ partnership, who have played outcome $o$, obtaining the stage game payoff $U_o$.

Let $\mathcal{P}_{h,o}^{ij\epsilon}$ be the (history-to-outcome) probability that an $ij$ partnership with history $h$ plays outcome $o$, which is:

- $(1-\epsilon)^2$ if $o$ is the outcome prescribed by strategies $i$ and $j$ given $h$.
- $\epsilon(1-\epsilon)$ if $o$ and the outcome prescribed by strategies $i$ and $j$ given $h$ differ only in the action of one player.
- $\epsilon^2$ if $o$ and the outcome prescribed by strategies $i$ and $j$ given $h$ differ in the action of both players.

At a steady state, the relationship between the distribution of $ij$ partnerships with history $h$ and those with resulting outcome $o$ is

$$w_o^{ij\epsilon} = \sum_{h \in H} z_h^{ij\epsilon} \mathcal{P}_{h,o}^{ij\epsilon}$$

or, in matrix form,

$$\boldsymbol{w}^{ij\epsilon} = \boldsymbol{z}^{ij\epsilon} \mathcal{P}^{ij\epsilon} \quad [1]$$

where $\boldsymbol{w}^{ij\epsilon}$ is a $1 \times 4$ row vector, $\boldsymbol{z}^{ij\epsilon}$ is a $1 \times 5$ row vector and $\mathcal{P}^{ij\epsilon}$ is a $5 \times 4$ matrix.

After playing the stage game, a partnership survives exogenous separation with probability $\delta$. Let $L_o^{ij} = 1$ if an $ij$ partnership is endogenously broken after outcome $o$, and $L_o^{ij} = 0$ otherwise. Consider a player involved in a $ij$ partnership where the outcome of the game is $o$ at a certain period. Then,

- With probability $\delta(1 - L_o^{ij})$, the partnership survives and the player's history becomes $h = o$ for the next period.
- With probability $(1-\delta) + \delta L_o^{ij}$, the partnership is broken and the player's history becomes $h = \emptyset$ for the next period.

Considering this, we obtain the outcome-to-history probabilities $N_{o,h}^{ij}$ (see table S2), which indicate the probability that an individual in an $ij$ partnership with outcome $o \in O$ (row heading in table S2) has the history $h \in H$ at the beginning of the next period (column heading in table S2).

At a steady state, the relationship between the distribution of $ij$ partnerships with outcome $o$ and those with history $h$ is then

$$z_h^{ij\epsilon} = \sum_{o \in O} w_o^{ij\epsilon} N_{o,h}^{ij}$$

or, in matrix form,

$$\boldsymbol{z}^{ij\epsilon} = \boldsymbol{w}^{ij\epsilon} N^{ij} \quad [2]$$

Combining Eq. (2) and Eq. (1) we obtain

$$\boldsymbol{z}^{ij\epsilon} = \boldsymbol{z}^{ij\epsilon}(\mathcal{P}^{ij\epsilon} N^{ij}) \quad [3]$$

and

$$\boldsymbol{w}^{ij\epsilon} = \boldsymbol{w}^{ij\epsilon}(N^{ij} \mathcal{P}^{ij\epsilon}) \quad [4]$$

Let $\boldsymbol{z}^{ij\epsilon*}$ be the distribution that solves Eq. (3) and let $\boldsymbol{w}^{ij\epsilon*}$ be the distribution that solves Eq. (4). For $\epsilon > 0$ these distributions exist and are unique, because $(\mathcal{P}^{ij\epsilon} N^{ij})$ and $(N^{ij} \mathcal{P}^{ij\epsilon})$ are (right or row) stochastic matrices with strictly positive entries (6).

Taking into account that $w_o^{ij\epsilon*}$ is the fraction of $i$-players (in $ij$ partnerships) who obtain payoff $U_o$, the average payoff obtained by $i$-players in $ij$ partnerships is

$$F_{ij}^{\epsilon} \equiv \sum_{o \in O} w_o^{ij\epsilon*} U_o$$

Let us normalize to 1 the total mass of players in the pool of singles (this is made to avoid introducing a new variable, it has no effect in the results). Then, the mass of $i$-players in new $ij$ partnerships each period is $p_i p_j$, and the total mass of $i$-players in (new and old) $ij$ partnerships is $m_{ij} \equiv \frac{p_i p_j}{z_\emptyset^{ij\epsilon*}}$, which implies

$$x_i = \frac{\sum_{j \in \mathbb{S}(\boldsymbol{x})} m_{ij}}{\sum_{k,j \in \mathbb{S}(\boldsymbol{x})} m_{kj}} = p_i \frac{\sum_{j \in \mathbb{S}(\boldsymbol{x})} \frac{p_j}{z_\emptyset^{ij\epsilon*}}}{\sum_{k,j \in \mathbb{S}(\boldsymbol{x})} \frac{p_k p_j}{z_\emptyset^{kj\epsilon*}}} \tag{5}$$

Equation Eq. (5) defines a function $f : \Delta_\Omega \to \Delta_\Omega$ such that $\boldsymbol{x} = f(\boldsymbol{p})$. For a population distribution $\boldsymbol{x}$, there is a unique pool distribution $\boldsymbol{p}^* = f^{-1}(\boldsymbol{x})$ such that $\boldsymbol{x} = f(\boldsymbol{p}^*)$ (see Proposition 1 in (7)), and $f^{-1}(\boldsymbol{x})$ can be computed using equation Eq. (5).

The average payoff obtained by $i$-players at pool distribution $\boldsymbol{p}$ is

$$\hat{F}_i^{\epsilon}(\boldsymbol{p}) = \frac{\sum_{j \in \mathbb{S}(\boldsymbol{x})} m_{ij} F_{ij}^{\epsilon}}{\sum_{j \in \mathbb{S}(\boldsymbol{x})} m_{ij}} = \frac{\sum_{j \in \mathbb{S}(\boldsymbol{x})} \frac{p_j}{z_\emptyset^{ij\epsilon*}} F_{ij}^{\epsilon}}{\sum_{j \in \mathbb{S}(\boldsymbol{x})} \frac{p_j}{z_\emptyset^{ij\epsilon*}}} \tag{6}$$

And the expected payoff to strategy $i$ at population $\boldsymbol{x}$ can consequently be calculated as

$$F_i^{\epsilon}(\boldsymbol{x}) = \hat{F}_i^{\epsilon}(f^{-1}(\boldsymbol{x}))$$

Monomorphic populations where every player uses strategy $j$ are represented as $\boldsymbol{e}_j$, and $F_{ij}^{\epsilon} = F_i^{\epsilon}(\boldsymbol{e}_j)$ is the payoff obtained by (a player using) strategy $i$ entering a monomorphic population of $j$-players.

**D. Strict Nash and evolutionary stable strategies.** In this section we study whether there are any strict Nash or evolutionarily stable strategies in our model. For the sake of clarity, we review these concepts below. Considering a strategy set $\Omega$:

- A (pure) strategy $i \in \Omega$ is a Nash equilibrium strategy if $F_{ii}^{\epsilon} \geq F_{ji}^{\epsilon}$ for every $j \in \Omega$. If the inequality is strict (i.e., $i$ is the unique best response to itself in $\Omega$), then $i$ is a **strict Nash strategy**.

- Besides being Nash, **a necessary condition for a strategy $i$ to be evolutionarily stable (ESS)** is

$$F_{ji}^{\epsilon} = F_{ii}^{\epsilon} \implies F_{ij}^{\epsilon} > F_{jj}^{\epsilon}$$

  for every $j \in \Omega$.

Evolutionary stability is a weaker condition than strict Nash, and both imply asymptotic stability under many evolutionary dynamics (with no experimentation). However, dynamic stability is not guaranteed if there is mutation or experimentation.

In the RPD with errors, we can find several strict Nash strategies (8, 9). For instance, using the techniques developed by (10), it can be shown that *AllD* is strict Nash in the RPD for any $\epsilon \in (0, 0.5)$ (see proof in section H). However, this is not the case in the VRPD. Focusing on the donation game, the following proposition shows that no memory-one strategy is evolutionarily stable (or strict Nash) in the VRPD in the type of settings we are interested in (i.e., small to moderate error level $\epsilon \in (0, 0.1]$ and $\delta \in [0.5, 1)$). The proof of this proposition is included in section H.

**Proposition 1.** *No memory-one strategy is evolutionarily stable in any donation game with endogenous separation, for any probability of continuation $\delta \in [0.5, 1)$ and any error rate $\epsilon \in (0, 0.1]$, not even considering only the restricted strategy space of memory-one strategies.*

Thus, in the RPD with occasional errors there are a number of monomorphic strict Nash states to which many evolutionary dynamics could converge, but this property is not kept in the VRPD. As an example, in our baseline scenario, *Grim*, *WSLS*, *AllD*, and $D|CDDC$ (*nWSLS*, for *nasty WSLS*) are strict Nash in the RPD (see proof in section H), but they are not in the VRPD (not even evolutionarily stable).

**E. Deterministic approximation: the mean dynamic.** In this section we derive the mean dynamic of our model. The mean dynamic (11, chapter 10; 12, chapter V) is a deterministic system of ODEs that approximates the (stochastic) dynamics of the agent-based model over finite time spans. If the population size is large enough and the revision probability is sufficiently low, the mean dynamic usually provides an excellent approximation to the stochastic dynamics. The mean dynamic of our model (described in the body of the paper) is the following differential equation*

$$\dot{x}_i = (1-\mu)\, \frac{x_i\, F_i^{\epsilon}(\boldsymbol{x})}{\bar{F}^{\epsilon}(\boldsymbol{x})} + \mu\, \frac{1}{n} - x_i \tag{7}$$

*For analytical convenience, to derive the mean dynamic, we have assumed that revising agents who experiment choose a feasible strategy at random, i.e. the chosen strategy may not be a neighbor of their current strategy, as in the stochastic model.

- $\dot{x}_i$ describes how the fraction of $i$-strategists evolves over time. There is one equation for each strategy $i \in \{1, ..., n\}$, with $n$ being the number of considered strategies. We can consider small subsets of strategies to observe how they fare against each other.

- $F_i^\epsilon(\boldsymbol{x})$ is the payoff obtained by strategy $i$ at population state $\boldsymbol{x}$ (see section C), and $\bar{F}^\epsilon(\boldsymbol{x}) \equiv \sum_{j=1}^{n} x_j F_j^\epsilon(\boldsymbol{x})$ is the average payoff in the population at state $\boldsymbol{x}$. The stage game payoffs are assumed to be non-negative (positive for symmetric action profiles), so the average payoff is always positive.

- The term $\frac{x_i\, F_i^\epsilon(\boldsymbol{x})}{\bar{F}^\epsilon(\boldsymbol{x})}$ corresponds to an inflow of agents using strategy $i$ that is proportional to the payoff obtained by each $i$-player $F_i^\epsilon(\boldsymbol{x})$ and to the proportion of $i$-players in the population. This term can be interpreted as the probability that an agent who revises its strategy adopts strategy $i$, or as the fraction of newborns that are expected to be of type $i$.

- The term $\mu\,\frac{1}{n}$ corresponds to the probability that a reviser experiments and adopts strategy $i$. It can also be interpreted as the probability that a newborn mutates and adopts strategy $i$.

- The last term $(-x_i)$ corresponds to the outflow of $i$-players (i.e., $i$-players who decide to revise their strategy or, under the biological interpretation, those who die), which is proportional to their presence in the population.

Equation (7) can equivalently be written as

$$\dot{x}_i = (1-\mu)\, x_i \frac{F_i^\epsilon(\boldsymbol{x}) - \bar{F}^\epsilon(\boldsymbol{x})}{\bar{F}^\epsilon(\boldsymbol{x})} + \mu\,(\frac{1}{n} - x_i), \tag{8}$$

which shows that the vector field (the right hand side of Eq. (7)) is a weighted composition of the Maynard-Smith replicator dynamics,

$$\dot{x}_i = x_i \frac{F_i^\epsilon(\boldsymbol{x}) - \bar{F}^\epsilon(\boldsymbol{x})}{\bar{F}^\epsilon(\boldsymbol{x})} \tag{9}$$

plus an experimentation field that introduces some uniform variability at every state. Equation (8) is a replication-mutation equation (13–16) with errors.

The stability of a rest point can be assessed by studying the eigenvalues of the Jacobian of the vector field at the rest point. If all the eigenvalues have negative real part, the rest point is asymptotically stable. At a rest point of Eq. (8), the eigenvalues $\lambda_\mu$ for Eq. (8) are related to the eigenvalues $\lambda$ for Eq. (9) by

$$\lambda_\mu = (1-\mu)\lambda - \mu$$

Assuming that the eigenvalues for Eq. (9) are bounded in the population space, it follows (17) that, for sufficiently large experimentation rate $\mu$, every rest point of Eq. (8) is asymptotically stable. And eventually, if the experimentation rate is large enough, there is a unique rest point of Eq. (8), which is asymptotically stable.

**F. The dominance of leavers.** In this section we solve the mean dynamic (MD) of our model numerically to better understand the dominance of strategies who exercise the option to leave. The *Mathematica* Notebook used to solve this MD is included in the accompanying repository. We start by exploring the MD of a system where the strategy space is composed of the three most successful strategies in the RPD (i.e., *Grim*, *WSLS* and *TFT*) plus $Leaver_1$. Figure S4 shows the solution trajectories starting from each of the four monomorphic states, with parameter values corresponding to the baseline scenario.

The four trajectories converge to the same state: 78% $Leaver_1$, 5% *Grim*, 12% *WSLS* and 5% *TFT*. We numerically solved the MD starting from 1000 random initial conditions and all the trajectories converged to the indicated state. This suggests that this state is a global attractor of the dynamics. The results for $Leaver_2$ and $Leaver_3$ are qualitatively the same.

We focus the discussion on $Leaver_1$; the arguments for the other *Leavers* are similar. Note that the sequence of outcomes when two players using $Leaver_1$ meet, assuming no exogenous breakup and no mistakes, is *DD-CC-CC-...* . A player's unilateral mistake would lead to endogenous separation, while a double mistake would lead to continuation with mutual cooperation.

With moderate levels of errors and experimentation, strategy $Leaver_1$ outperforms *Grim*, *WSLS* and *TFT* in bimorphic populations, as shown in the bottom row of figure S5. Each plot in figure S5 includes the (1-dimensional) MD at the bottom of each plot: arrows point in the direction of movement and circles indicate the location of rest points (black if asymptotically stable, or white if unstable). The accompanying repository includes an interactive *Mathematica* notebook that can be used to replicate all these figures and analyze any bimorphic population.

Panels a1 (no errors, no experimentation) and a2 (errors and experimentation) compare $Leaver_1$ with *Grim*. Note that errors affect the expected payoff obtained by each strategy (the payoff functions), while experimentation does not affect the payoff functions but affects the dynamics, including the existence and location of rest points. Comparing panels a1 and a2, it is clear that the payoff obtained by $Leaver_1$ (solid line) is very robust to errors, while *Grim* suffers a considerable loss. Two *Grim* players turn to mutual defection after a mistake, and do not restore cooperation unless there is a double mistake; in contrast, $Leaver_1$ players break up after a mistake and can go back to mutual cooperation with a new $Leaver_1$ partner after one period. With the error rate of the baseline scenario (panel a2), the payoff obtained by *Grim* is always below the payoff obtained by $Leaver_1$.

Panels b1 and b2 compare $Leaver_1$ with *WSLS*. In this case, the payoff differences are smaller, but $Leaver_1$ is slightly more robust to errors than *WSLS*, and, under the error rate considered in the baseline scenario (panel b2), $Leaver_1$ obtains slightly larger payoffs.

Panels c1 and c2 compare $Leaver_1$ with *TFT*. Again, $Leaver_1$ (solid line) is more robust to errors than *TFT* (dashed line). Panel c2 shows that, when the proportion of *TFT* players is very large (left part of the panel), *TFT* obtains a larger payoff than $Leaver_1$, but the small basin of attraction that should correspond to *TFT* is wiped out by experimentation. Note on the phase portrait at the bottom of panel c2 that, due to experimentation: i) the strict Nash state where every player uses *TFT* is not even a rest point, ii) there is no rest point either at the internal Nash state where the payoffs are equal, and iii) there is a single attractor where most players (but not all) use $Leaver_1$.

To wrap up this section, let us explore the performance of the three *Leavers* in a population initially dominated by the four most successful strategies in the model without the option to leave (i.e., *Grim*, *WSLS*, *TFT* and $C|CDCC$). Figure S6 shows a representative solution trajectory, which converges to a state where the three *Leavers* dominate. We have numerically solved this MD starting from 1000 random initial conditions and all the trajectories converged to the same state. This suggests that this system has a unique global attractor, where more than 85% of players are using one of the three *Leaver* strategies, while each of the four strategies that do not leave accounts for less than 5% of the population (in a system where the experimentation rate is already 5%).

Figure S7 shows how the strategy distribution of the seemingly unique global attractor of this MD changes with the experimentation rate $\mu$. As the experimentation rate $\mu$ tends to zero, the three *Leaver* strategies increase their dominance and end up completely eliminating the four strategies that do not leave.

**G. Small errors vs. infinitesimally small errors.** Figure S8 has been created to provide a tentative explanation of the sharp change in the distribution of strategies shown in figure 6 of the paper. In figure S8, for very low error rates ($\epsilon \leq 10^{-3}$) there is a unique global attractor where the presence of $Leaver_1$ is very low. Then, there is a small region of values of $\epsilon$ for which there are two attractors: the previous one and a new one with much greater presence of $Leaver_1$ (see $\epsilon = 4 \times 10^{-3}$). And finally, for $\epsilon \geq 5 \times 10^{-3}$, the second attractor becomes global. This new global attractor is characterized by the dominance of $Leaver_1$, and stays for greater values of $\epsilon$.

#### H. Proofs.

*Proof of Proposition 1.* Let $\Omega$ be the set of 162 memory-one strategies, and let $F^{\epsilon}_{ij}$ be the payoff to strategy $i$ in a population of $j$-players, with noise $\epsilon$ (section C). The proof consists in showing that, for every strategy $i \in \Omega$, and for $\delta$ and $\epsilon$ in the considered ranges, either (a) there is another strategy $j \in \Omega$ such that $F^{\epsilon}_{ji} = F^{\epsilon}_{ii} = F^{\epsilon}_{ij} = F^{\epsilon}_{jj}$ (so $i$ is not ESS), or (b) there is another strategy $j \in \Omega$ with $F^{\epsilon}_{ji} > F^{\epsilon}_{ii}$ (so $i$ is not even a Nash equilibrium strategy).

Let us define the set $\underline{\Omega}$ as the set of all memory-one strategies $\Omega$ except for $C|DDDD$, $C|DCDD$, $C|DDCD$, $D|CCCC$, $D|CCDC$ and $D|CDCC$. For each outcome $o \in \{CC, CD, DC, DD\}$ let $o^*$ be the outcome $o$ seen from the perspective of the other player; e.g., if $o = CD$, then $o^* = DC$, while if $o = CC$, then $o^* = o = CC$. Let $i(h) \in \{C, D, L\}$ be strategy $i$'s prescribed action after history $h \in \{\emptyset, CC, CD, DC, DD\}$. Note that, for every strategy $i \in \underline{\Omega}$, there is either

- some outcome $o \in \{CC, CD, DC, DD\}$ after which strategy $i$ leaves, i.e., such that

$$i(o) = L \qquad [10]$$

- or, otherwise, some outcome $o$ that makes both players in an $ii$ partnership adopt the initial action $i(\emptyset)$ again, i.e., such that

$$i(o) = i(o^*) = i(\emptyset) \qquad [11]$$

This opens the door for the existence of another strategy $j \in \Omega$ that generates the same sequence of outcomes as strategy $i$ in contests in which only strategies $i$ and $j$ are involved, so $F^{\epsilon}_{ji} = F^{\epsilon}_{ii} = F^{\epsilon}_{ij} = F^{\epsilon}_{jj}$. We prove this by construction.

To build $j \in \Omega$ from $i \in \underline{\Omega}$, let $o$ be an outcome satisfying Eq. (10) or, if $i$ never leaves, satisfying Eq. (11).

- If strategy $i \in \underline{\Omega}$ leaves after outcome $o$, i.e., $i(o) = L$, then define $j \in \Omega$ exactly as $i$ but replacing the action after outcome $o^*$, $j(o^*)$, with:
  - If $o = o^*$, take $j(o^*) = j(o) = i(\emptyset)$. In this case, any $ii$ or $ij$ partnership breaks up after outcome $o$, and any $jj$ partnership goes on playing as if the partnership had been broken.
  - If $o \neq o^*$, take $j(o^*) \neq i(o^*)$. In this case, any $ii$, $ij$ or $jj$ partnership breaks up after players observe $o/o^*$ or $o^*/o$, because $i(o) = j(o) = L$.
- If strategy $i \in \underline{\Omega}$ never leaves and there is an outcome $o$ such that $i(o) = i(o^*) = i(\emptyset)$, then define $j \in \Omega$ exactly as $i$ but with $j(o^*) = L$. In this case, any $jj$ partnership breaks up after observing $o/o^*$, any $ii$ partnership goes on playing as if the partnership had been broken, and any $ij$ partnership may do one or the other (depending on whether $o = o^*$ and, if not, on which player observes $o$ and which observes $o^*$).

With these conditions, the expected sequence of outcomes for an $i$-player and a $j$-player in a population made up by $i$-players and $j$-players is the same (after the same outcome, some partnership combinations may break up while others stay together, but the next outcome follows the same probability distribution), so $F^{\epsilon}_{ji} = F^{\epsilon}_{ii} = F^{\epsilon}_{ij} = F^{\epsilon}_{jj}$.

Up to now we have shown that no strategy in $\underline{\Omega}$ is evolutionarily stable, for any symmetric 2x2 stage game, any $\delta$ and any $\epsilon$. Focusing now on the donation game, it only remains to show that none of the other six memory-one strategies is evolutionarily stable.[†]

- Taking $i = D|CDCC$ and $j = D|CDDC$, the calculation of $F^{\epsilon}_{ji} - F^{\epsilon}_{ii}$ for the donation game leads to an expression that is positive for $\delta(1-2\epsilon)^2 \geq \epsilon > 0$. If this condition is satisfied, then $D|CDCC$ is not an equilibrium strategy. For instance, for $0 < \epsilon \leq 0.1$, the condition is satisfied if $\delta > \frac{0.1}{0.8^2} \approx 0.16$. For $0 < \epsilon \leq 0.15$, the condition is satisfied if $\delta > \frac{0.15}{0.7^2} \approx 0.31$. If $\epsilon = 0$, then $F^{\epsilon}_{ji} = F^{\epsilon}_{ii} = F^{\epsilon}_{ij} = F^{\epsilon}_{jj}$.
- Taking $i \in \{C|DDDD, C|DCDD, C|DDCD\}$ and $j = D|LLLL$,we find that $F^{\epsilon}_{ji} - F^{\epsilon}_{ii}$ for the donation game is positive for $0 \leq \epsilon < 0.5$ and $0 < \delta < 1$. Consequently, none of the considered three strategies (with initial action $C$) are equilibrium strategies.
- Taking $i \in \{D|CCCC, D|CCDC\}$ and $j = D|DDDD$, we find that $F^{\epsilon}_{ji} - F^{\epsilon}_{ii}$ for the donation game is positive for $0 \leq \epsilon < 0.5$ and $0 < \delta < 1$. Consequently, $D|CCCC$ and $D|CCDC$ are not equilibrium strategies.

□

*Proof of Statement:* ***AllD* is strict Nash in the RPD, for any $\epsilon \in (0, 0.5)$.**

*AllD* can be implemented as a state space strategy (9) with one single state $S_D$ where action $D$ is chosen, no matter the partnership's actual history of outcomes. Consequently, if the stage payoff obtained by one of the actions (C or D, with errors) against *AllD* is greater than the other, the action with the largest stage payoff is optimal at every stage. The stage payoff for $C$ against *AllD* is

$$(1-\epsilon)^2 U_{CD} + \epsilon(1-\epsilon)(U_{CC} + U_{DD}) + \epsilon^2 U_{DC}$$

The stage payoff for $D$ against *AllD* is

$$(1-\epsilon)^2 U_{DD} + \epsilon(1-\epsilon)(U_{DC} + U_{CD}) + \epsilon^2 U_{CC}$$

For the Prisoner's Dilemma and $0 \leq \epsilon < 0.5$, the payoff for $D$ is greater than the payoff for $C$: the payoff difference is a second degree polynomial in $\epsilon$

$$f(\epsilon) = (U_{DD} - U_{CD}) + \epsilon(U_{DC} - U_{CC} - 3U_{DD} + 3U_{CD}) + 2\epsilon^2(U_{DD} - U_{CD} - U_{DC} + U_{CC})$$

with $f(0) = U_{DD} - U_{CD} > 0$, $f(0.5) = 0$ and no interior minimum for $\epsilon \in (0, 0.5)$. For $\epsilon > 0$, every possible history of outcomes may happen with positive probability. Consequently, for $0 < \epsilon < 0.5$ the unique best response to *AllD* is the strategy that maps every history to action $D$. That strategy is precisely *AllD* itself. □

*Proof of Statement:* **In our baseline scenario, *Grim*, *WSLS*, *AllD*, and *nWSLS* are strict Nash in the RPD.**

For *AllD* see the previous proof. We only detail the proof for *Grim*, as the method for the other two strategies is similar.

In a stationary population of $i$-players and $j$-players with no endogenous separation, the mass of $i$-players in $t$-period-old $ij$ partnerships, $m^{[t]}_{ij}$ satisfies $m^{[t+1]}_{ij} = \delta m^{[t]}_{ij}$. Adding the payoff obtained by all $i$-players in $ij$ partnerships (of any longevity) and dividing by their mass $\frac{m^{[1]}_{ij}}{1-\delta}$ we obtain that the expected payoff to an $i$-player in a population of $j$-players is

$$F^{\epsilon}_{ij} = (1-\delta) \sum_{t=1}^{\infty} \delta^{t-1} \sum_{o \in \{C,D\}^2} w^{\epsilon,t}_{ij}(o) U_o \qquad [12]$$

where:

- $w^{\epsilon,t}_{ij}(o)$ is the fraction of (ordered) $o$-outcomes at the $t^{th}$ repetition of the stage game, in $ij$ partnerships.
- $U_o$ is the stage game payoff (to the $i$-player) for outcome $o$.

Equation (12) turns out to be equal to the standard normalized payoff for repeated games with no separation, where $\delta$ is interpreted as a discount factor. We can consequently use dynamic programming techniques as in the standard case of repeated games with no separation (9).

*Grim* can be implemented as a state space strategy with two states (9, 10): $S_D$ (where action $D$ is chosen) and $S_C$ (where action $C$ is chosen, and initial state). Outcome $CC$ leads to state $S_C$, and all other outcomes lead to $S_D$. Let:

- $(1-\delta)V_{S_C}$ be the expected payoff obtained by a strategy that plays optimally against *Grim*, when facing *Grim* in state $S_C$.

[†] The accompanying repository contains a *Mathematica* notebook named *proposition-1-proof-assistant.nb* that can be used to check this statement.

- $(1-\delta)V_{S_D}$ be the expected payoff obtained by a strategy that plays optimally against *Grim*, when facing *Grim* in state $S_D$.

- $(1-\delta)V_{C,S_C}$ be the expected payoff obtained by a strategy that plays $C$ and, afterwards, optimally against *Grim*, when facing *Grim* in state $S_C$. The definitions for $V_{D,S_C}$, $V_{D,S_C}$ and $V_{D,S_C}$ are equivalent. From these definitions and Eq. (12) we have:

$$
\begin{aligned}
V_{C,S_C} &= (1-\epsilon)^2(U_{CC}+\delta V_{S_C}) + \epsilon(1-\epsilon)(U_{DC}+U_{CD}+2\delta V_{S_D}) + \epsilon^2(U_{DD}+\delta V_{S_D}) \\
V_{D,S_C} &= (1-\epsilon)^2(U_{DC}+\delta V_{S_D}) + \epsilon(1-\epsilon)(U_{CC}+\delta V_{S_C}+U_{DD}+\delta V_{S_D}) + \epsilon^2(U_{CD}+\delta V_{S_D}) \\
V_{C,S_D} &= (1-\epsilon)^2(U_{CD}+\delta V_{S_D}) + \epsilon(1-\epsilon)(U_{CC}+\delta V_{S_C}+U_{DD}+\delta V_{S_D}) + \epsilon^2(U_{DC}+\delta V_{S_D}) \\
V_{D,S_D} &= (1-\epsilon)^2(U_{DD}+\delta V_{S_D}) + \epsilon(1-\epsilon)(U_{DC}+U_{CD}+2\delta V_{S_D}) + \epsilon^2(U_{CC}+\delta V_{S_C})
\end{aligned} \quad [13]
$$

with $V_{S_C} = max(V_{C,S_C}, V_{D,S_C})$ and $V_{S_D} = max(V_{C,S_D}, V_{D,S_D})$. These equations have a unique solution (9).

*Grim* is a strict Nash strategy if the action prescribed by *Grim* at each state is the unique optimal action, that is, if

$$V_{C,S_C} > V_{D,S_C} \text{ and } V_{D,S_D} > V_{C,S_D}, \quad [14]$$

in which case $V_{S_C} = V_{C,S_C}$ and $V_{S_D} = V_{D,S_D}$, leading to the system of equations:

$$
\begin{aligned}
V_{S_C} &= (1-\epsilon)^2(U_{CC}+\delta V_{S_C}) + \epsilon(1-\epsilon)(U_{DC}+U_{CD}+2\delta V_{S_D}) + \epsilon^2(U_{DD}+\delta V_{S_D}) \\
V_{S_D} &= (1-\epsilon)^2(U_{DD}+\delta V_{S_D}) + \epsilon(1-\epsilon)(U_{DC}+U_{CD}+2\delta V_{S_D}) + \epsilon^2(U_{CC}+\delta V_{S_C})
\end{aligned} \quad [15]
$$

Solving Eq. (15) for our baseline scenario provides $V_{S_C} = 35.00$ and $V_{S_D} = 22.59$ , and then, from Eq. (13), $V_{C,S_C} = 35.00 > V_{D,S_C} = 25.82$, $V_{D,S_D} = 22.59 > V_{C,S_D} = 22.22$, so Eq. (14) holds. □

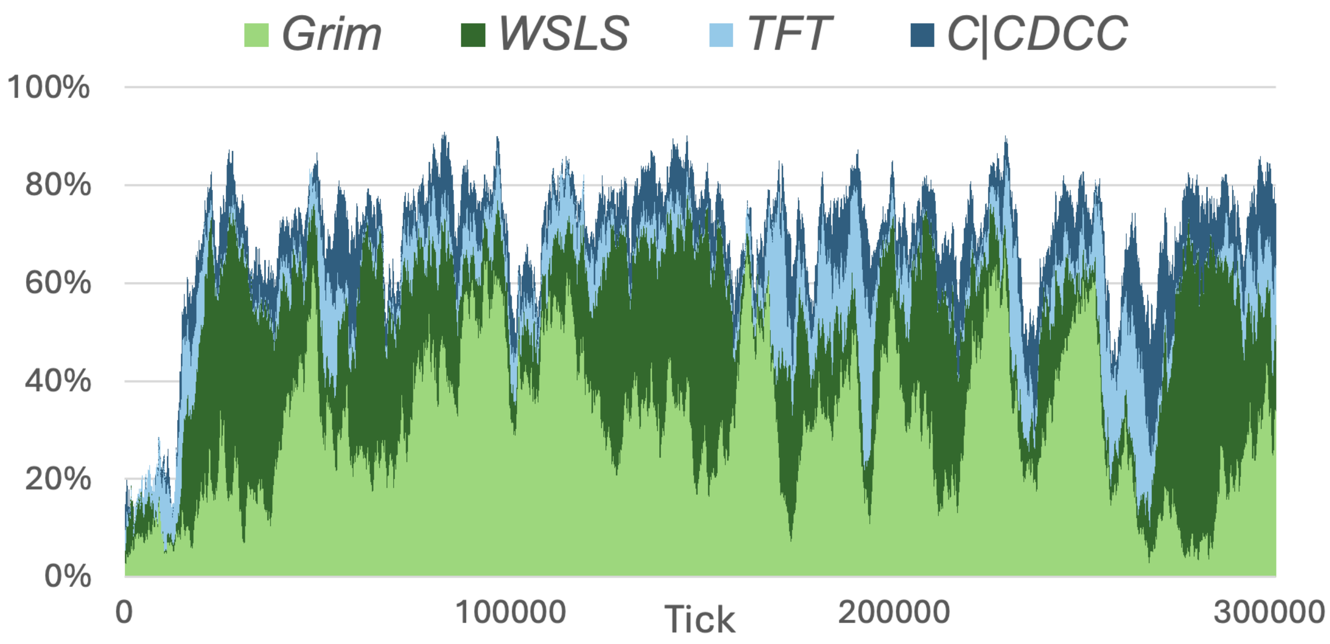


**Fig. S1.** Percentage of the population using each of the four main strategies, plotted against time, in a representative run of the RPD in the baseline scenario. It is a stacked line graph, so the frequency of *Grim* is the height of the light green line, the frequency of *WSLS* is the distance between the dark green line and the light green line, and so on.

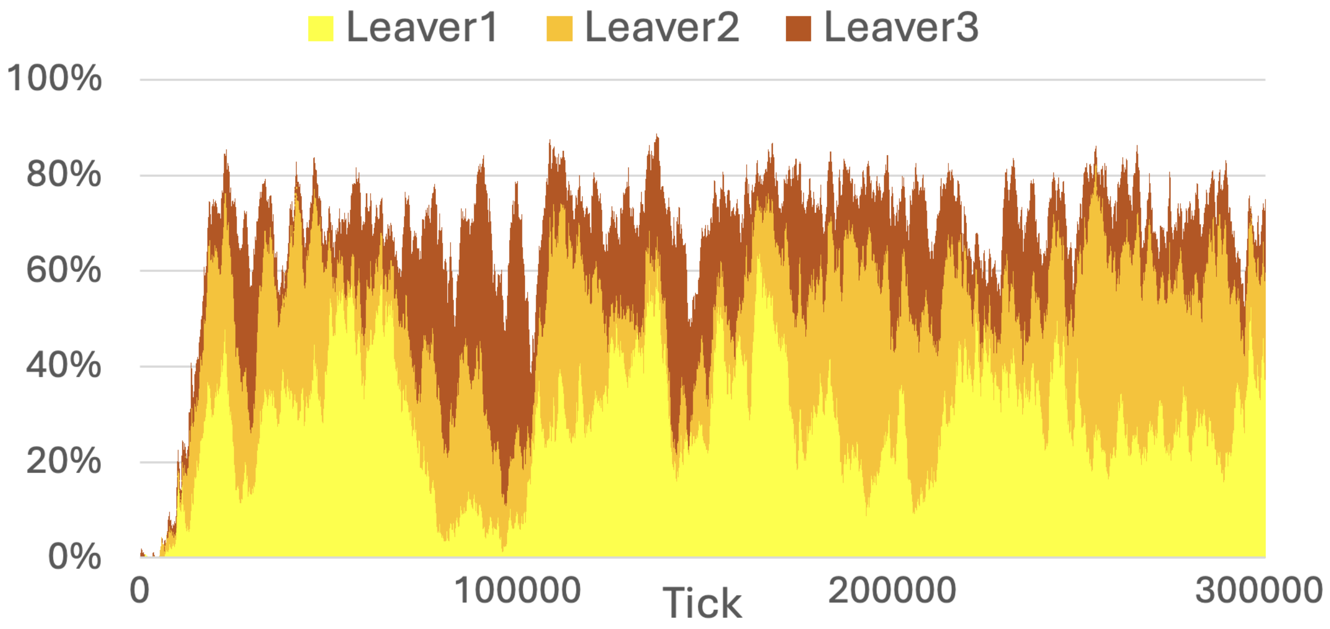


**Fig. S2.** Percentage of the population using each of the three main strategies plotted against time, in a representative run of the model with the option to leave in the baseline scenario.It is a stacked line graph, so that the frequency of Leaver 1 is the height of the yellow line, the frequency of Leaver 2 is the distance between the orange line and the yellow line and so on.

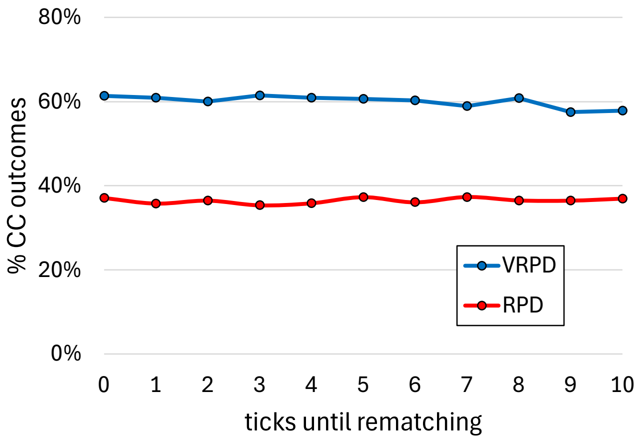


**Fig. S3.** Average percentage of cooperative outcomes in the RPD and VRPD, departing from the baseline scenario, for different values of parameter *ticks-until-rematching* and with *out-payoff* equal to 0. Each point is the average from tick 190 000 to tick 200 000 over 100 runs starting from random initial conditions. The standard error of every point is below $2\%$. The baseline scenario corresponds to *ticks-until-rematching* = 0.

**Table S1. Percentage of players using each choice (row) after a given history (column) in the VRPD, for three different values of *ticks-until-rematching*: 0, 5 and 10. Values gathered as in Table 2 of the paper.**

| | | History | | | | | | | | | | | | | | |
|---|---|---|---|---|---|---|---|---|---|---|---|---|---|---|---|---|
| | | $\emptyset$ | | | $CC$ | | | $CD$ | | | $DC$ | | | $DD$ | | |
| Ticks until rematching | | 0 | 5 | 10 | 0 | 5 | 10 | 0 | 5 | 10 | 0 | 5 | 10 | 0 | 5 | 10 |
| | C | 3 | 5 | 7 | **89** | **82** | **74** | 4 | 3 | 2 | 26 | 23 | 22 | **94** | **90** | **85** |
| Choice | D | **97** | **95** | **93** | 9 | 14 | 20 | 15 | 13 | 11 | 46 | 46 | 47 | 5 | 7 | 9 |
| | L | | | | 2 | 4 | 6 | **82** | **84** | **87** | 28 | 31 | 31 | 1 | 3 | 6 |

**Table S2. Probability that a player in a partnership with outcome $o$ (row) starts the following period with history $h \in H$ (column).**

| | | History $h$ | | | | |
|---|---|---|---|---|---|---|
| | | $\emptyset$ | $CC$ | $CD$ | $DC$ | $DD$ |
| $o$ | $CC$ | $(1-\delta)+\delta L^{ij}_{CC}$ | $\delta(1-L^{ij}_{CC})$ | 0 | 0 | 0 |
| | $CD$ | $(1-\delta)+\delta L^{ij}_{CD}$ | 0 | $\delta(1-L^{ij}_{CD})$ | 0 | 0 |
| | $DC$ | $(1-\delta)+\delta L^{ij}_{DC}$ | 0 | 0 | $\delta(1-L^{ij}_{DC})$ | 0 |
| | $DD$ | $(1-\delta)+\delta L^{ij}_{DD}$ | 0 | 0 | 0 | $\delta(1-L^{ij}_{DD})$ |

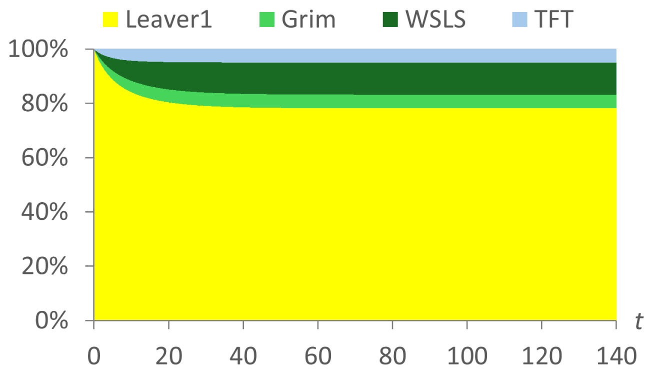


**(a)** Starting from a population of $Leaver_1$

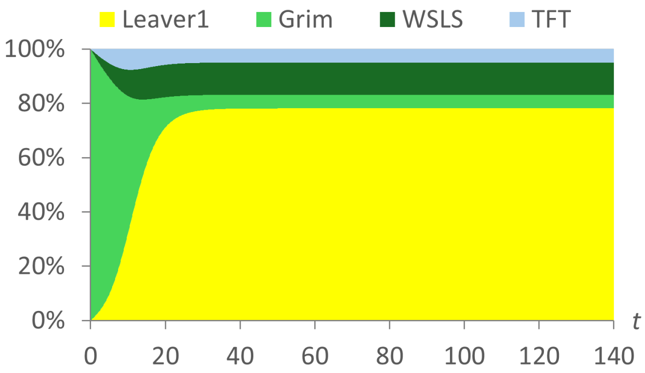


**(b)** Starting from a population of *Grim*

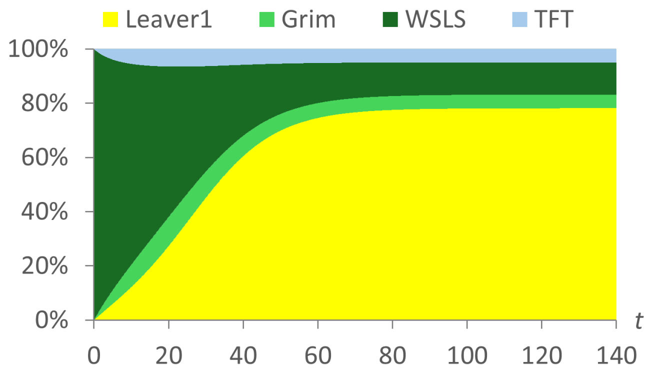


**(c)** Starting from a population of *WSLS*

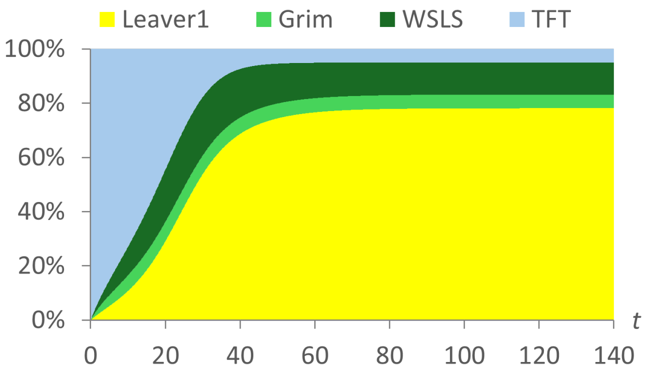


**(d)** Starting from a population of *TFT*

**Fig. S4.** Solution trajectories of the mean dynamic starting from a monomorphic population of (a) $Leaver_1$, (b) *Grim*, (c) *WSLS*, and (d) *TFT*. Parameter values are set as in the baseline scenario.

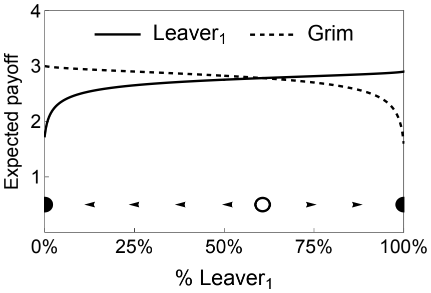


(a1) $Leaver_1$ vs *Grim*; $\epsilon = \mu = 0$

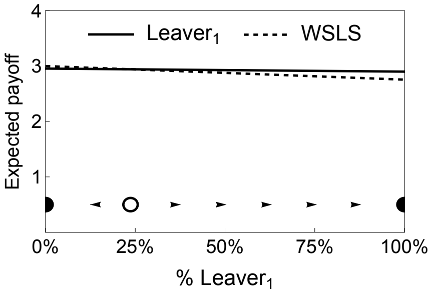


(b1) $Leaver_1$ vs *WSLS*; $\epsilon = \mu = 0$

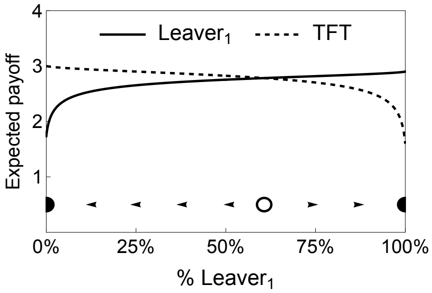


(c1) $Leaver_1$ vs *TFT*; $\epsilon = \mu = 0$

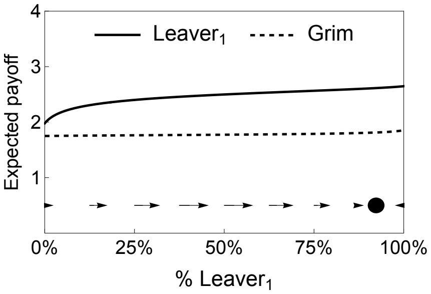


(a2) $Leaver_1$ vs *Grim*; $\epsilon = \mu = 0.05$

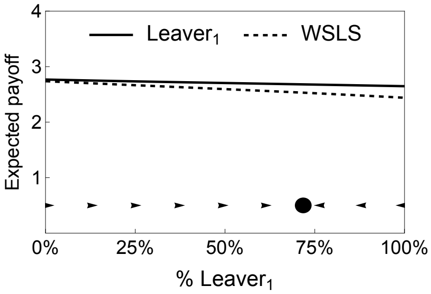


(b2) $Leaver_1$ vs *WSLS*; $\epsilon = \mu = 0.05$

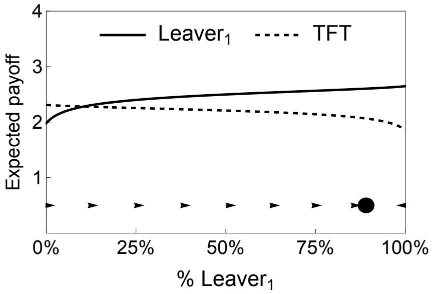


(c2) $Leaver_1$ vs *TFT*; $\epsilon = \mu = 0.05$

**Fig. S5.** Payoff functions and phase portraits for several bimorphic populations in the VRPD. In a $\{Leaver_1, Grim\}$ population (subfigures a1 and a2), *Grim* suffers more with errors, and the dynamics basically wipe out *Grim*. In a $\{Leaver_1, WSLS\}$ population (subfigures b1 and b2), *WSLS* is almost as robust to errors as $Leaver_1$, although the presence of errors gives a small advantage to $Leaver_1$. In a $\{Leaver_1, TFT\}$ population (subfigures c1 and c2), *TFT* suffers more with errors, and it is basically wiped out. Parameters are set as in the baseline scenario except where indicated otherwise.

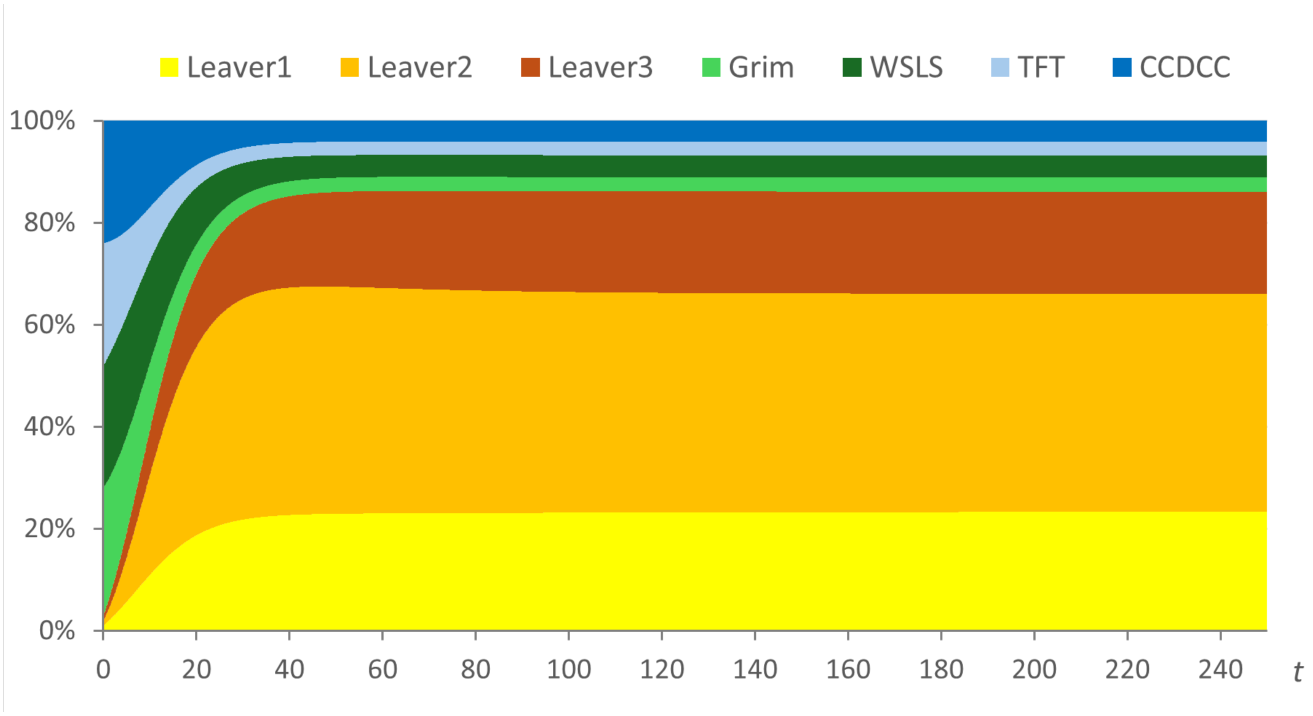


**Fig. S6.** Solution trajectory of the MD of a system that includes strategies *Grim*, *WSLS*, *TFT* and $C|CDCC$, together with the three $Leaver$ strategies. Parameter values are set as in the baseline scenario.

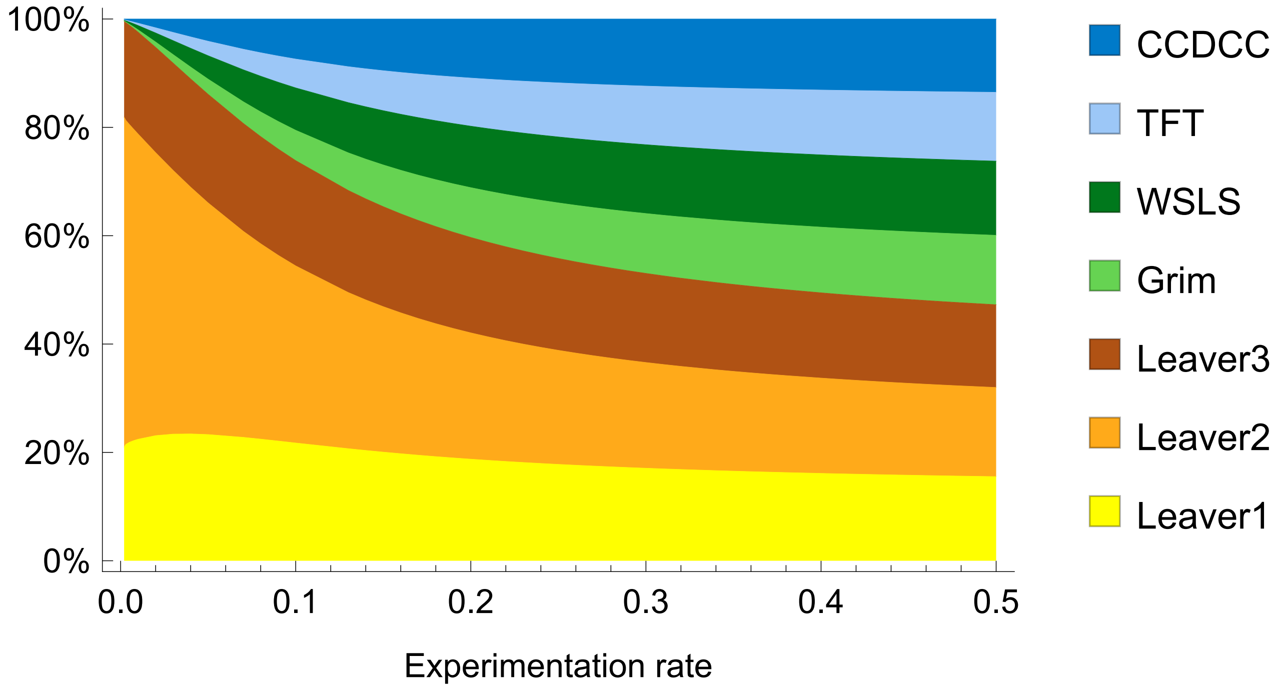


**Fig. S7.** Strategy distribution of the seemingly unique global attractor of the MD for different values of the experimentation rate $\mu \in (0, 0.5]$. Other parameter values are set as in the baseline scenario.

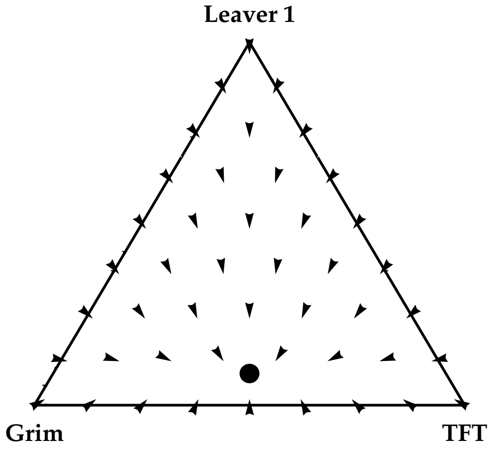

**(a)** Error rate $\epsilon = 0$

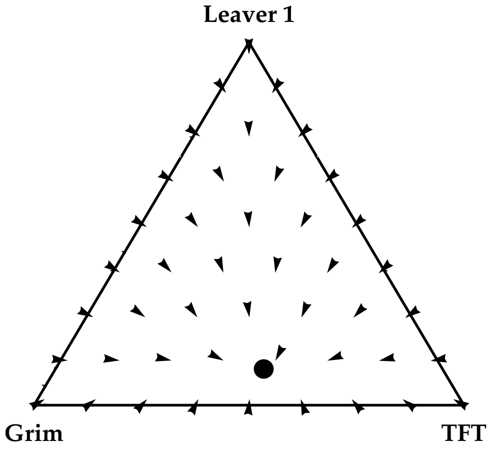

**(b)** Error rate $\epsilon = 10^{-3}$

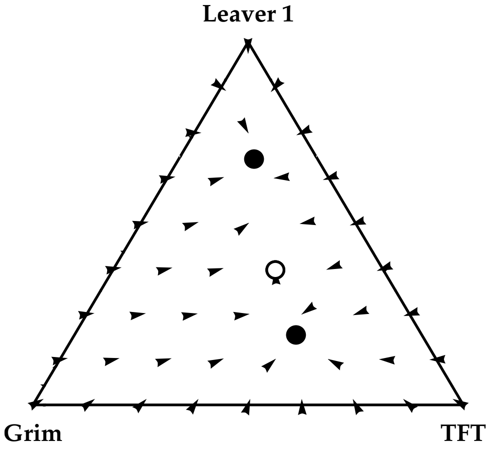

**(c)** Error rate $\epsilon = 4 \times 10^{-3}$

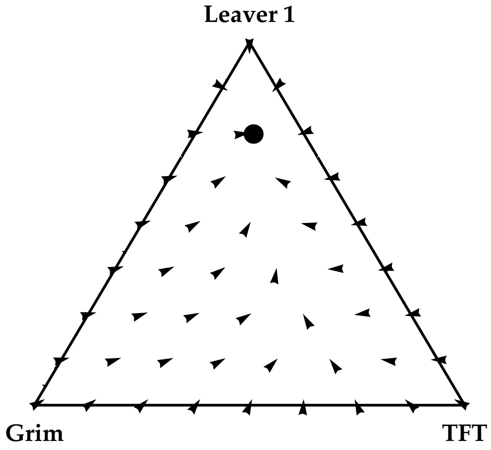

**(d)** Error rate $\epsilon = 5 \times 10^{-3}$

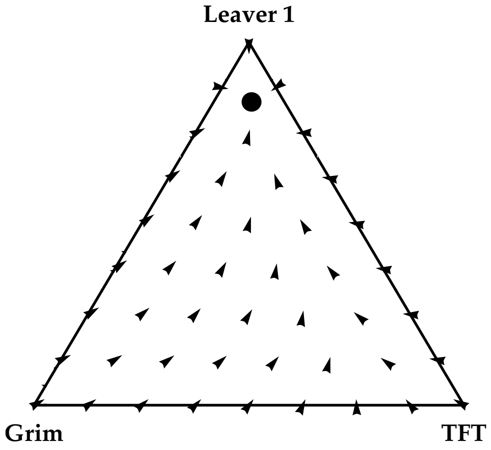

**(e)** Error rate $\epsilon = 0.01$

**Fig. S8.** Mean dynamic of the system with strategies $Leaver_1$, *Grim* and *TFT*, for different error rates $\epsilon$. Isolated rest points are represented with circles: black if the rest point is asymptotically stable, white if unstable. Other parameter values are set as in the baseline scenario.

### References


1. CF Chabris, JJ Lee, D Cesarini, DJ Benjamin, DI Laibson, The fourth law of behavior genetics. *Curr. directions psychological science* **24**, 304–312 (2015).
2. H Kokko, R Brooks, JM McNamara, AI Houston, The sexual selection continuum. *Proc. Royal Soc. London. Ser. B: Biol. Sci.* **269**, 1331–1340 (2002).
3. A Traulsen, D Semmann, RD Sommerfeld, HJ Krambeck, M Milinski, Human strategy updating in evolutionary games. *Proc. Natl. Acad. Sci.* **107**, 2962–2966 (2010).
4. J Grujić, et al., A comparative analysis of spatial prisoner's dilemma experiments: Conditional cooperation and payoff irrelevance. *Sci. Reports* **4**, 4615 (2014).
5. J Tkadlec, C Hilbe, MA Nowak, Mutation enhances cooperation in direct reciprocity. *Proc. Natl. Acad. Sci.* **120**, e2221080120 (2023).
6. RA Horn, CR Johnson, *Matrix Analysis.* (Cambridge University Press), (1985).
7. SS Izquierdo, LR Izquierdo, Stable strategies in repeated games with endogenous separation. *Universidad de Valladolid Discuss. Pap.* (2026).
8. YG Kim, Evolutionarily stable strategies in the repeated prisoner's dilemma. *Math. Soc. Sci.* **28**, 167–197 (1994).
9. O Leimar, Repeated games: A state space approach. *J. Theor. Biol.* **184**, 471–498 (1997).
10. R Boyd, Mistakes allow evolutionary stability in the repeated prisoner's dilemma game. *J. Theor. Biol.* **136**, 47–56 (1989).
11. WH Sandholm, *Population games and evolutionary dynamics.* (The MIT Press), (2010).
12. LR Izquierdo, SS Izquierdo, WH Sandholm, *Agent-Based Evolutionary Game Dynamics.* (University of Wisconsin Pressbooks), (2024).
13. J Hofbauer, The selection mutation equation. *J. Math. Biol.* **23**, 41–53 (1985).
14. P Stadler, P Schuster, Mutation in autocatalytic reaction networks. *J. Math. Biol.* **30**, 597–632 (1992).
15. KM Page, MA Nowak, Unifying evolutionary dynamics. *J. Theor. Biol.* **219**, 93–98 (2002).
16. SS Izquierdo, LR Izquierdo, Strictly dominated strategies in the replicator-mutator dynamics. *Games* **2**, 355–364 (2011).
17. J Bauer, M Broom, E Alonso, The stabilization of equilibria in evolutionary game dynamics through mutation: mutation limits in evolutionary games. *Proc. Royal Soc. A: Math. Phys. Eng. Sci.* **475**, 20190355 (2019).
